# MOD: A novel machine-learning optimal-filtering method for accurate and efficient detection of subthreshold synaptic events *in vivo*

**DOI:** 10.1101/2020.07.04.186478

**Authors:** Xiaomin Zhang, Alois Schlögl, David Vandael, Peter Jonas

**Affiliations:** IST Austria (Institute of Science and Technology Austria), Am Campus 1, A-3400 Klosterneuburg, Austria

**Author notes:** Equal contribution (Xiaomin Zhang, Alois Schlögl). **Corresponding author:** Dr. Peter Jonas, IST Austria, Am Campus 1, A-3400 Klosterneuburg, Austria.

**Keywords:** Subthreshold activity, excitatory postsynaptic potentials (EPSPs), excitatory postsynaptic currents (EPSCs), *in vivo* recordings, supervised machine learning, Wiener filter, cross-validation, single-neuron computations, neuronal coding

## Abstract

To understand the mechanisms of information coding in single neurons, it is necessary to analyze subthreshold synaptic events, action potentials (APs), and the interrelation between these two forms of activity in different behavioral states. However, detecting excitatory postsynaptic potentials (EPSPs) or currents (EPSCs) in awake, behaving animals remains challenging, because of unfavorable signal-to-noise ratio, high frequency, fluctuating amplitude, and variable time course of synaptic events. Here, we developed a new method for synaptic event detection, termed MOD (**M**achine-learning **O**ptimal-filtering **D**etection-procedure), which combines concepts of supervised machine learning and optimal Wiener filtering. First, experts were asked to manually score short epochs of data. Second, the algorithm was trained to obtain the optimal filter coefficients of a Wiener filter and the optimal detection threshold. Third, scored and unscored data were processed with the optimal filter, and events were detected as peaks above threshold. Finally, the area under the curve (AUC) of the receiver operating characteristics (ROC) curve was used to quantify accuracy and efficiency of detection. Additionally, cross-validation was performed to exclude overfitting of the scored data, a potential concern with machine-learning approaches. We then challenged the new detection method with EPSP traces *in vivo* in mice during spatial navigation and EPSC traces *in vitro* in slices under conditions of enhanced transmitter release. When benchmarked using a (1−AUC)^−1^ metric, MOD outperformed previous methods (template-fit and deconvolution) by a factor of up to 3. Thus, MOD may become an important tool for large-scale analysis of synaptic activity *in vivo* and *in vitro*.

**Highlights:**

- A new method for detection of synaptic events, termed MOD, is described
- The method combines the concepts of supervised machine learning and optimal filtering
- The method is useful for analysis of both *in vitro* and *in vivo* data sets
- MOD outperforms previously published methods for synaptic event detection by a factor of up to 3

## Introduction

To fully understand the nature of the neural code, it is necessary to analyze the rules of single-neuron computations, by which neurons convert analogue synaptic input signals into digital synaptic output signals (Koch, 1999; Debanne et al., 2013). Ideally, one would like to simultaneously probe subthreshold excitatory postsynaptic potentials (EPSPs) and suprathreshold action potentials (APs) *in vivo* in defined behavioral conditions. Whereas huge progress has been made on the experimental side, notably in the fields of *in vivo* patch-clamp recording (Lee et al., 2009; Pernía-Andrade and Jonas, 2014), *in vivo* extracellular measurements (Stark et al., 2012; Jun et al., 2017), and *in vivo* optogenetics (Scanziani and Häusser, 2009; Deisseroth, 2015), the development of techniques for adequate analysis has lagged behind. The main difficulties are: (1) the low signal-to-noise ratio, as synaptic events are often related to the activity of a small number of synapses, (2) the high-frequency of synaptic events, leading to a high degree of temporal overlap, (3) the slow time course of the synaptic events, which often are recorded as EPSPs rather than excitatory postsynaptic currents (EPSCs), because of lack of sufficient voltage-clamp conditions, and (4) the variability in synaptic amplitude and kinetics, as events are often generated by synapses located on different dendritic compartments. Thus, reliable and efficient detection of synaptic activity in *in vivo* data sets remains challenging.

For the simpler but related problem of detection of spontaneous synaptic activity *in vitro* (Kavalali, 2015), several different methods were proposed. These include amplitude threshold methods, derivative-based methods (Maier et al., 2011), template-fit algorithms (Jonas et al., 1993; Clements and Bekkers, 1997; Chadderton et al., 2004), deconvolution methods (Pernía-Andrade et al., 2012), and Bayesian approaches that consider distributions of templates rather than single templates (Merel et al., 2016). However, these methods cannot be easily applied to *in vivo* data sets. In such conditions, none of the techniques reaches the sensitivity, reliability, and temporal resolution of manual analysis by an experienced expert, who has prior knowledge of the time course of the synaptic events and can relatively easily distinguish true events from experimental artifacts. On the other hand, manual scoring of large *in vivo* data sets is not practicable, because of the extremely time consuming nature of the analysis. Recently, machine learning-based techniques have been used to infer spiking activity from measured Ca^2+^ transients (Sasaki et al., 2008; Theis et al., 2016; Berens et al., 2018). However, whether machine learning-based strategies can be exploited for the detection of subthreshold synaptic events remains unclear. Furthermore, machine learning approaches are often computationally demanding, which may be a limiting factor for large-scale analysis of synaptic data.

To analyze subthreshold synaptic activity *in vivo* with improved accuracy and efficiency, we developed a new detection method, rooted in the concepts of supervised machine learning (Kevin and Murphy, 2012) and optimal filtering (Wiener and Hopf, 1931). First, short epochs of EPSP or EPSC data were manually scored by experts. Second, the detection algorithm was trained to predict the expert scoring. During training, an optimal set of filter coefficients was computed using the Wiener-Hopf equation. Finally, the approach was tested by cross-validation, splitting the scored data into various combinations of training and test sets. When benchmarked on experimentally recorded data sets, the new method greatly outperformed previous methods, especially for *in vivo*, but also for *in vitro* data sets.

## Methods

### Detection of synaptic events by machine learning and optimal filtering

A number of detection methods for spontaneous synaptic events were previously proposed, including template fit (Jonas et al., 1993; Clements and Bekkers, 1997; Chadderton et al., 2004) and deconvolution (Pernía-Andrade et al., 2012). Although these methods work well in *in vitro* measurements and *in vivo* in anesthetized animals, it is difficult to apply them to *in vivo* recordings from awake, behaving animals. The main reasons are unfavorable signal-to-noise ratio, high frequency, and variable amplitude and time course of synaptic events. Thus, the detection performance of all available methods is substantially below that of a trained expert. To overcome these limitations, we developed a novel approach combining the detection power of machine learning with the computational efficiency of optimal filtering. In previous methods, the algorithm detects putative events, which are individually validated by an expert user (Jonas et al., 1993; Clements and Bekkers, 1997; Chadderton et al., 2004; Pernía-Andrade et al., 2012). In the new approach, the expert first manually scores a short stretch of data. This information is then used to train a detection algorithm, which is why the method is conceptually related to “machine learning”. Based on the training, an optimal filter is generated, which is able to convert input traces into output traces (“raw detection traces”) that closely resemble the manual scoring traces. To efficiently compute the optimal filter coefficients, we use the Wiener-Hopf equations (Wiener and Hopf, 1931, Wiener, 1949), which is why the method is related to “optimal filtering” or “Wiener filtering”. Finally, we apply the optimal filter to scored and unscored data, enabling validation of the method and large-scale, high-throughput analysis of subthreshold activity *in vivo* and *in vitro*. The new technique was termed MOD, which stands for **M**achine-learning and **O**ptimal-filtering-based **D**etection-procedure.

### Details of implementation of Machine-learning Optimal-filtering Detection-procedure (MOD) Expert scoring

To define synaptic events and distinguish them from baseline noise, two experts (one with *in vivo* recording and analysis background and one with *in vitro* recording and analysis background) independently scored a subset of data (∼30 s at the beginning (S1) and ∼30 s at the end of the recording (S2); **Table S1**). Events were manually scored in “SigViewer” (https://github.com/cbrnr/sigviewer; version 0.5.1), which allows us to set markers and annotations to the data and save them as separate event files. The experts were asked to put the event marker to a consistent fiducial point throughout the entire scoring period (e.g. onset or peak).

To account for possible jitter in marker positioning, each time point was symmetrically extended around the marker by a total window length *t*_win_. This generated a so-called manual scoring trace of zeros (0) and ones (1), with the same length and sampling frequency as the original data.

### Optimal Wiener filtering

The basic idea of MOD is to generate an optimal filter, which converts the original data into a raw detection trace closely resembling the manual scoring trace. As the data represented a discrete time series with a fixed sampling frequency, we used an optimal finite impulse response (FIR) filter, also known as “Wiener filter”. The transfer function of such a filter (G(z)) in the *z* domain is represented by

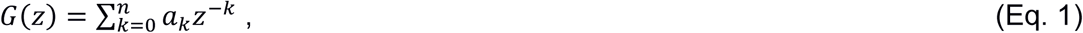

where *a*_0_, *a*_1_, …, *a*_*n*_ represent the filter coefficients, *n* + 1 is the order of the filter, and *z* is a complex number (Oppenheim and Schafer, 2010). The filter operation is then defined by

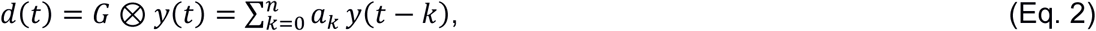

where *y(t)* denotes the recorded original data trace, *d(t)* is the raw detection trace, and *G* is the transfer function. The filter is considered causal, because it uses only sample points from “present” and “past” (i.e *y*(*t*), *y*(*t*−1),… *y*(*t*−*k*)), but no sample points from the “future”. For an optimal filter, the raw detection trace *d(t)* should resemble the manual scoring trace *s(t)* as closely as possible. Thus, we needed to find the optimum filter coefficients 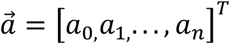 that minimized the sum of squared errors ∑_*t*_ *e*(t)^2^ with

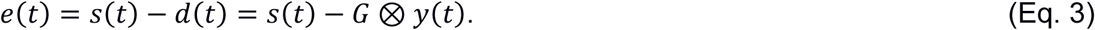

As the amplitude of the signal component of the detection trace is 1, minimizing *e*(*t*) is equivalent to maximizing the signal-to-noise ratio. Under the assumption of zero-mean processes *y*(*t*) and *s*(*t*), the filter coefficients can be computed by solving the Wiener-Hopf equation (Wiener and Hopf, 1931) as

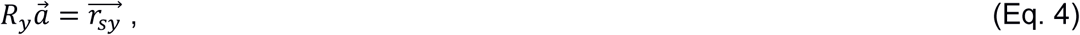

where *R*_*y*_ is a Toeplitz matrix of the auto-correlation function of the original data *y*(*t*) with

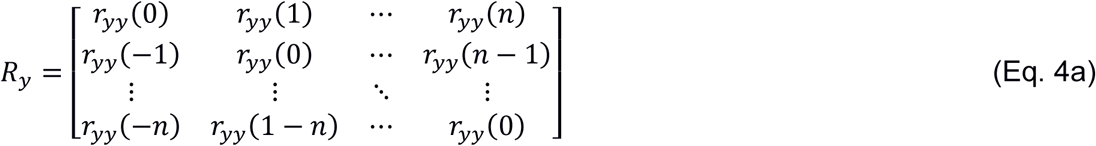

and 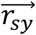 is a vector of the cross-correlation function between observed data *y*(*t*) and scoring trace *s*(*t*) with

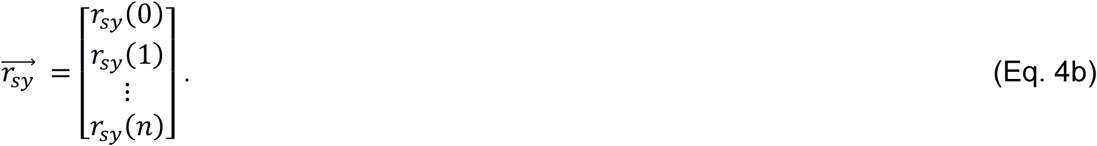

Then, the filter coefficients 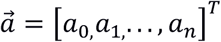 are obtained by solving the system of linear equations (Eq. 4) as

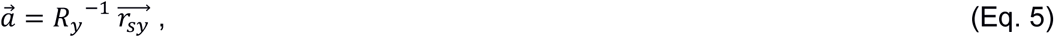

where *R*_y_^-1^ denotes the inverse of *R*_*y*_. Both auto- and cross-correlation functions were estimated from the samples of the training data set (*t* ∈ *trainSet*). In order to avoid any estimation bias caused by non-zero mean, the overall mean values µ_*y*_ and µ_*s*_ were removed from *y* and *s*, respectively. The auto-correlation function was computed as

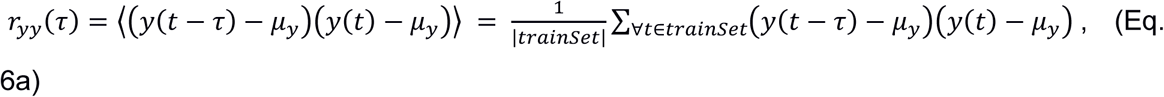

where ⟨*x*⟩ is the expectation value of *x*. Thus, *r*_*yy*_(*τ*) is symmetric (i.e. *r*_*yy*_(*τ*) = *r*_*yy*_(−*τ*)) and has its maximum at *τ* = 0. Exploiting its symmetry properties, the auto-correlation function needs to be computed only for lag values of 0 ≤ *τ* ≤ *n*. Moreover, the matrix *R*_*y*_ is positive definite (assuming the number of samples in the training set is sufficiently large, i.e. *N* >> *n*, where *N* is the total number of samples in the training set). The cross-correlation function is defined as

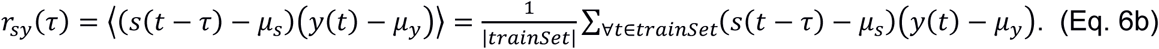

The standard implementation of Wiener filter is that of a causal filter that uses *r*_*yy*_(*τ*), *r*_*sy*_(*τ*) for *τ* ≥ 0. In practice, we determined the filter coefficients for a series of time-shifted cross-correlation functions from *r*′_*sy*_(*τ*) = *r*_*sy*_(*τ* − *δ*) for *τ* ≥ 0 in the Wiener-Hopf equations such that

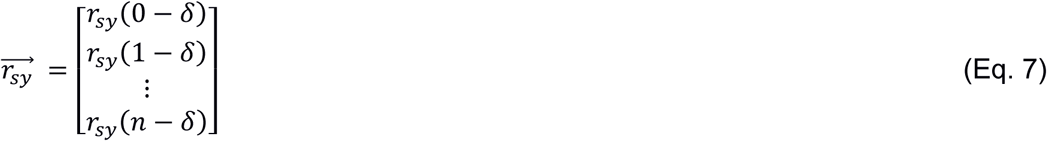

and we determined the detection trace according to the equation

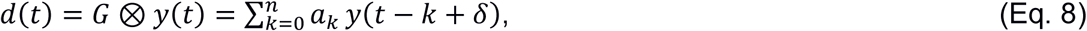

where *δ* represents a time shift. This implementation may introduce non-causal properties of the filter, because −*k* + *δ* can become > 0. In order to determine the optimum time shift, Wiener filtering was performed for various values of *δ* (*δ* = −10 ms to 40 ms, varied in steps of 0.2 ms). The value that produced the highest detection performance was consistently applied to the experimental data (see below; **Fig. S3**). Biosig 3.5.0 (http://biosig.sourceforge.net/) was used for loading the data and for the AUC, the ROC and κ analysis. The NaN-toolbox 3.1.4 (https://octave.sourceforge.io/nan/) was used for handling of missing or invalid data samples when computing the auto- and cross-correlation functions according to *IEEE 854-1987* standard. Finally, the raw detection trace was low-pass filtered in forward and reverse direction using a Hann window of order 13, corresponding to an effective cutoff frequency of 1.075 kHz.

### Detection performance

To quantify the performance of the detection algorithm, ROC curves were computed (Schlögl et al., 2007; Pernía-Andrade et al., 2012; Berens et al., 2018). These curves describe the relation between true positive rate (TPR) and false positive rate (FPR) for various levels of detection threshold. Next, the area under each ROC curve (AUC) was determined (**Fig. 2C, F; Fig. 3C**; **Fig. 4C**). We used a fast algorithm based on sorting the sample values of the detection trace, such that every possible detection threshold was represented as a point in the ROC curve. For reasonable detection algorithms, the AUC could vary between 0.5 and 1.0, with 1.0 corresponding to perfect detection and 0.5 representing randomness.

**Fig. 1.**
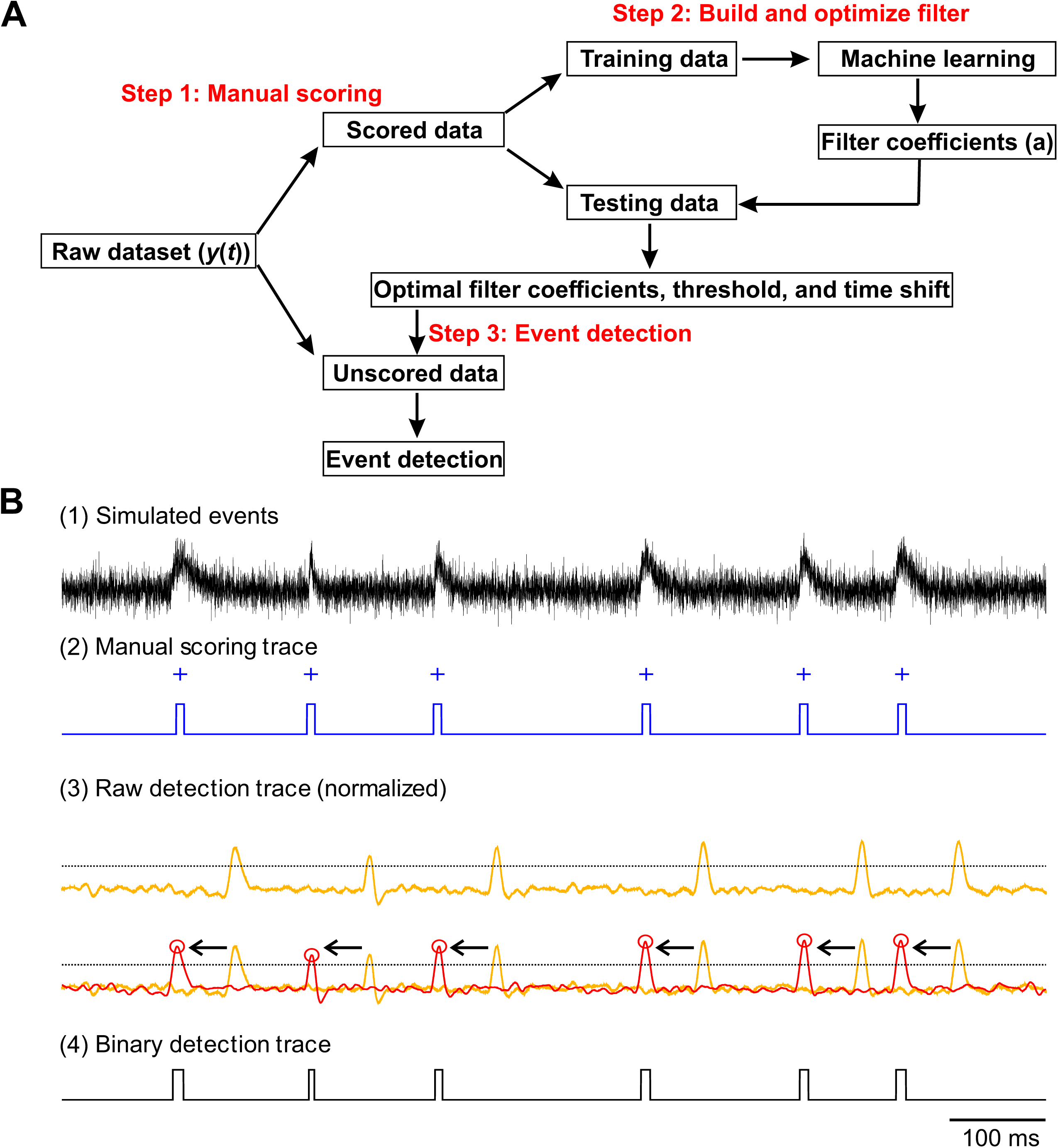
Flowchart of MOD analysis. **(A)** Flowchart of the MOD procedure. After the raw data was recorded, parts of the data sets were manually scored by experts. Using these manually scored events, the algorithm was then trained to produce an output that closely resembles the manual scoring trace. Optimal filter coefficients were then calculated based on the Wiener-Hopf equation, and the optimal filter was subsequently applied to the original data to generate a raw detection trace. **(B)** Input and output of the MOD procedure. Traces show, from top to bottom: (1) schematic original data trace, (2) manual scoring of individual synaptic events (scoring markers ‘+’) and manual scoring trace generated by applying a symmetric ± 2 ms window to each marker, (3) raw detection trace generated by the MOD algorithm (yellow, unshifted; red, shifted; circles represent maxima of shifted trace; dotted horizontal line indicates threshold computed according to the maximum of Cohen’s κ; arrows indicate time shift δ, which compensates for causal properties of the Wiener filter and systematic differences in marker positioning between experts), and (4) binary detection trace. Note that the raw detection trace shows a substantially improved signal-to-noise ratio.

**Fig. 2.**
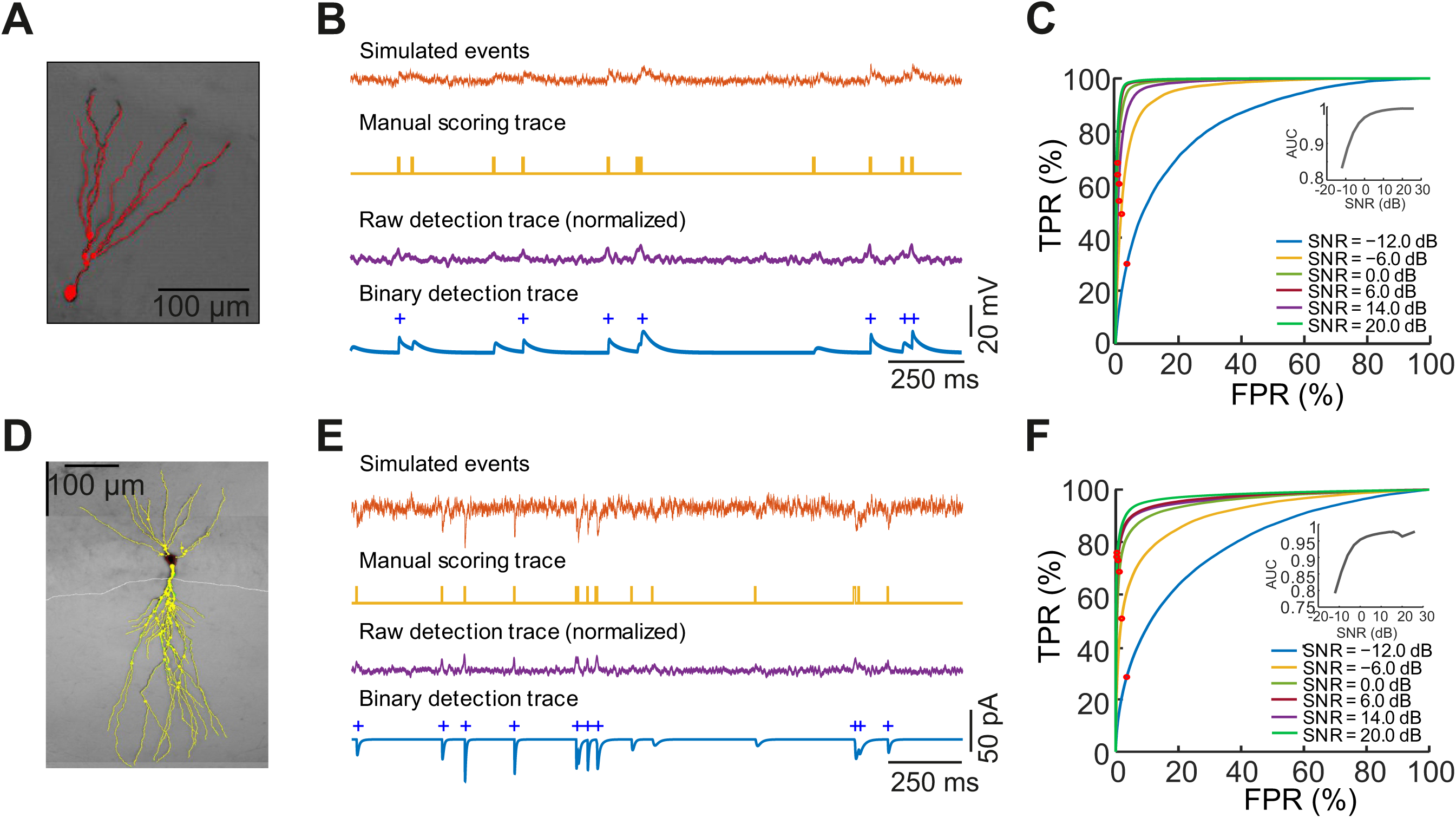
Test of MOD analysis on ground truth data obtained by cable model simulations. **(A)** Light micrograph of biocytin-labeled GC from *in vivo* recording data set. Reconstruction is superimposed in red. **(B)** Input and output of the MOD procedure. Traces show, from top to bottom: (1) simulated original trace, with signal-to-noise ratio of 0 dB (2) ground-truth scoring trace generated by applying a symmetric ± 2 ms window to each marker, (3) raw detection trace generated by the MOD algorithm, and (4) detected synaptic events (detection markers ‘+’), superimposed with the original data. **(C)** ROC curve analysis of detection performance of MOD applied to synthetic data sets. TPR was plotted against FPR. ROC curves for different signal-to-noise ratios between −12 and 20 dB. Noise was added according to a colored noise model. Note that the AUC is > 0.9 in many cases, demonstrating the accuracy and efficiency of the method. Inset, plot of AUC against signal-to-noise ratio. Note that the AUC remains high over a wide range of signal-to-noise ratios, even for signal-to-noise ratios < 1. (**D–F**) Similar plots as (A–C), but for *in vitro* EPSC data. Light micrograph of biocytin-labeled CA3 pyramidal neuron from *in vitro* recording data set. Reconstruction is superimposed in yellow.

**Fig. 3.**
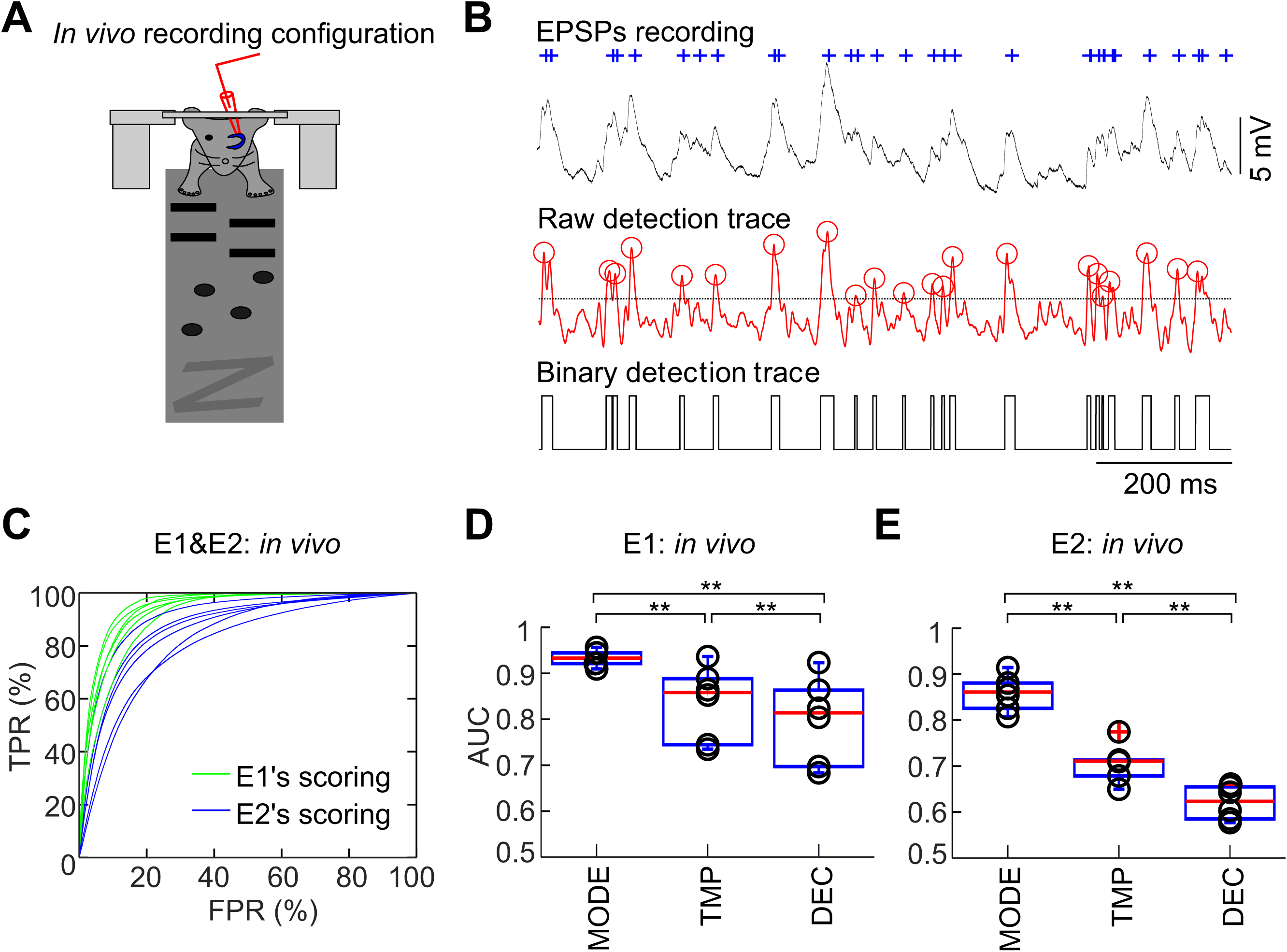
MOD permits efficient and accurate detection of EPSPs *in vivo*. **(A)** *In vivo* recording configuration. Whole-cell patch-clamp recording from dentate gyrus GCs in head-fixed animal running on a linear belt. The belt showed three partitions with different texture, providing spatial cues to the animal during the experiment. **(B)** A one-second segment of scored *in vivo* data. EPSPs were recorded under current-clamp conditions near the resting potential of the recorded cell. Top, raw data, together with the scoring from one expert. Center, raw detection trace from the MOD method; the markers (∘) indicate peaks above threshold. Horizontal dotted line indicates the optimum threshold corresponding to the maximum of Cohen’s κ. Bottom, binary detection trace obtained by applying the optimum threshold to the raw detection trace. **(C)** ROC curve analysis of detection performance. TPR was plotted against FPR. Data were from six *in vivo* data sets; each data set was scored by two independent experts E1 (green) and E2 (blue). Note that the AUC is close to 0.9, demonstrating the reliability and efficiency of the method. (**D, E**) Comparison of the performance of MOD against previously published methods (TMP, template fit; DEC, deconvolution). Summary bar graphs show the AUC values for six *in vivo* data sets based on two independent scorings (circles, individual data points; red line, median; box, interquartile range; whiskers, most extreme data points that are no more than 1.5 × IQR from box edge). Note that MOD outperforms the traditional methods TMP and DEC for EPSPs recorded *in vivo* (** indicates P < 0.01). Results shown were computed using the cross-validation scheme “A1B2–A2B1” with *t*_win_ = 4 ms.

**Fig. 4.**
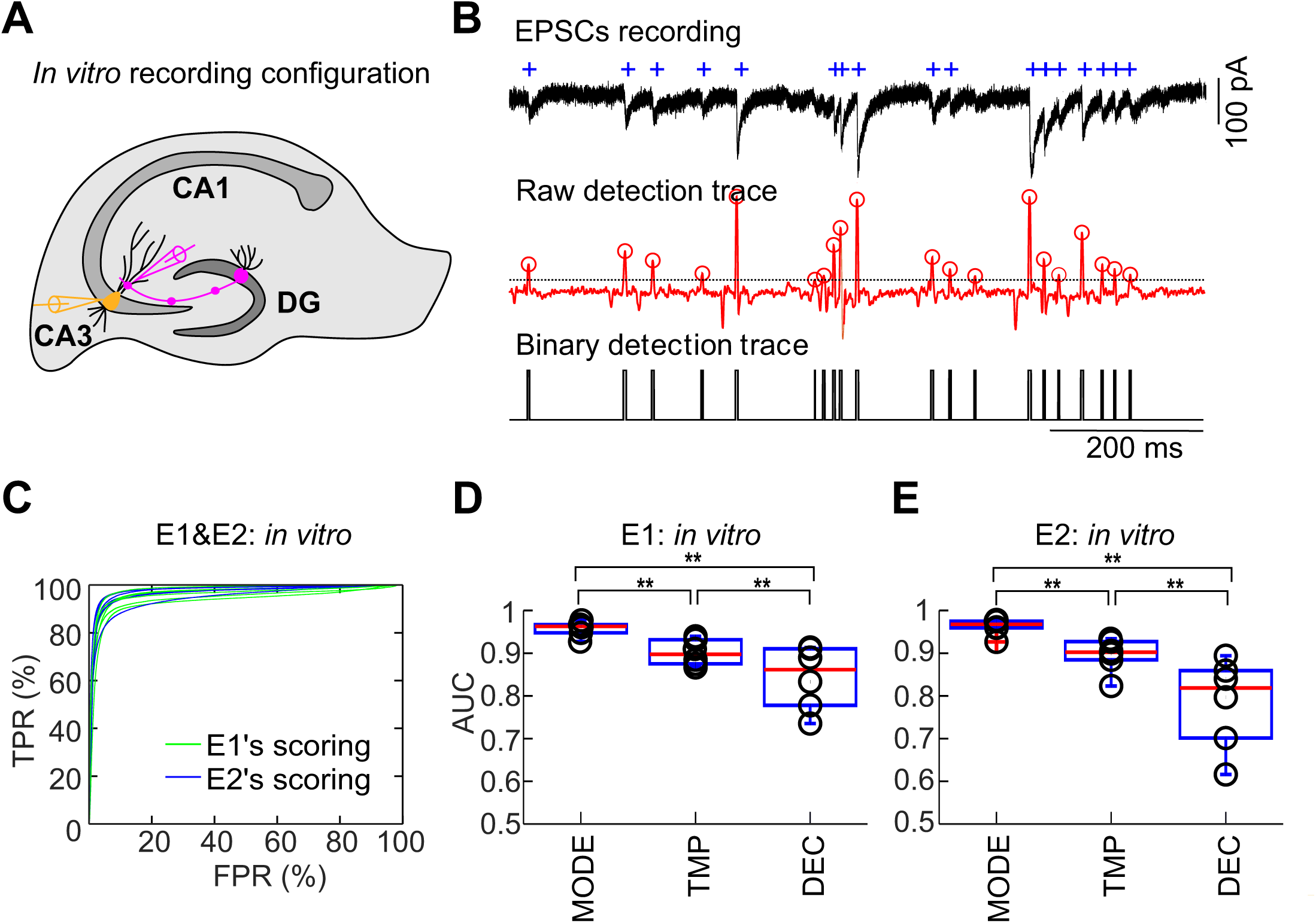
MOD permits efficient and accurate detection of EPSCs *in vitro*. **(A)** *In vitro* recording configuration. Cell-attached patch-clamp recording from a hippocampal mossy fiber terminal, combined with whole-cell patch-clamp recording from a postsynaptic CA3 pyramidal neuron in an acute brain slice. A 1-s, 100-Hz stimulus train was applied to the presynaptic terminal. **(B)** A one-second segment of scored *in vitro* data. EPSCs were recorded under voltage-clamp conditions at a holding potential of −70 mV. Top, raw data, together with the scoring from one expert. Center, raw detection trace from the MOD method, the markers (∘) indicate peaks above threshold. Horizontal dotted line indicates the optimum threshold corresponding to the maximum of Cohen’s κ. Bottom, binary detection trace obtained by applying the optimum threshold to the raw detection trace. **(C)** ROC curve analysis of detection performance. TPR was plotted against FPR. Data were from six *in vitro* data sets; each data set was scored by two independent experts E1 (green) and E2 (blue). Note that the AUC is close to 0.95. Thus, the performance of MOD approaches that of an ideal detector for the *in vitro* data set. (**D, E**) Comparison of the performance of MOD against previously published methods (TMP, template fit; DEC, deconvolution). Summary bar graphs show the AUC values for six *in vitro* data sets based on two independent scorings (circles, individual data points; red line, median; box, interquartile range; whiskers, most extreme data points that are no more than 1.5 × IQR from box edge). Note that MOD outperforms the traditional approaches TMP and DEC for EPSCs recorded *in vitro* (** indicates P < 0.01). Results shown were computed using the cross-validation scheme “A1B2–A2B1” with *t*_win_ = 4 ms.

In addition to allowing the benchmarking of different detection methods (**Fig. 3D, E**; **Fig. 4D, E**), AUC analysis permitted the optimization of free parameters in the analysis method, including the window *t*_win_ of the scoring trace (**Fig. S1**) and the time shift δ of the Wiener filter (**Fig. S3**). Finally, AUC analysis was used to probe the effect of additional preprocessing of the original data, such as low-pass and notch filtering. For further analysis, we used the values that produced the highest AUC values.

### Computing optimal detection threshold and binary detection trace

The Wiener filtering approach provided a raw detection trace with optimal signal-to-noise ratio; however, it did not determine the detection threshold. To find the optimal detection threshold (θ_opt_), Cohen’s κ coefficient was computed for every possible threshold (i.e. all points on the ROC curve). The threshold that maximized Cohen’s κ was then used to convert the raw detection trace into a binary detection trace (**Fig. 1B**).

Cohen’s κ was computed as:

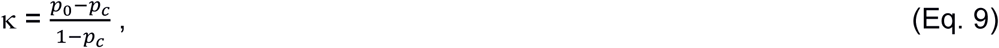

where p_0_ is the observed agreement between manual scoring trace and binary detection trace, and p_c_ is the agreement expected by chance (Cohen, 1960; Powers, 2012). Cohen’s κ was chosen over other measures because it was well bounded between −1 and 1. It showed clear maximum behavior as the threshold was shifted to higher values, and appeared to be particularly suitable for asymmetric problems, in which the absolute numbers of events in two classes were very different. Accordingly, several performance measures (including: mutual information, Youden index, accuracy, Matthews correlation coefficient, and F1 score) appeared to be less suitable (**Fig. S5B**), because either TPR was too low (e.g. accuracy) or FPR was too high (e.g. mutual information).

### Cross-validation

A possible pitfall in machine learning approaches is over-fitting of the data (Bishop, 2006). In an extreme scenario, it is possible to find a perfect classifier for a given data set, if the number of free parameters is comparable to the number of events. However, it is unclear if the classifier generalizes to new data. To rigorously test this, we used several schemes for cross-validation, which account for both within- and across-cell variation (**Fig. 5**). (1) Within-cell scheme (S1– S2): For each cell, S1 was used as training set and S2 as the test set. Subsequently, training and test sets were reversed. This cross-validation scheme tested whether the classification method generalized over the entire recording time. (2) Within-cell-split-half scheme (A1B2– A2B1): For each cell, scoring was performed at the beginning (S1) and at the end (S2) of the experimental recording time. Each section was then split into two halves, A and B. The training set was assembled from the first half of S1 and the second half of S2 (i.e. A1B2), and the test set was composed of the second half of S1 and the first half of S2 (A2B1). Subsequently, training and test sets were reversed. This cross-validation scheme tested whether the classification method generalized to adjacent new data. (3) Leave-one-(cell)-out-method (LOOM): In this scheme the data from all but one cell (the training set) were used to build the classifier, and this classifier was applied to compute the detection trace of the unused cell (the test set). The data sets were permuted until every set was included exactly once in the test set. This cross-validation method provided a benchmark of how well the detection performed on a new individual cell of the same type. Additionally, we performed cross-validation after adding additional pre-processing steps (200 and 1000 Hz low-pass filter, 1-Hz high-pass filter, 50-Hz notch filter). Differences in the AUC were < 1.46%, suggesting that preprocessing of the original data is neither necessary nor advantageous in the process.

**Fig. 5.**
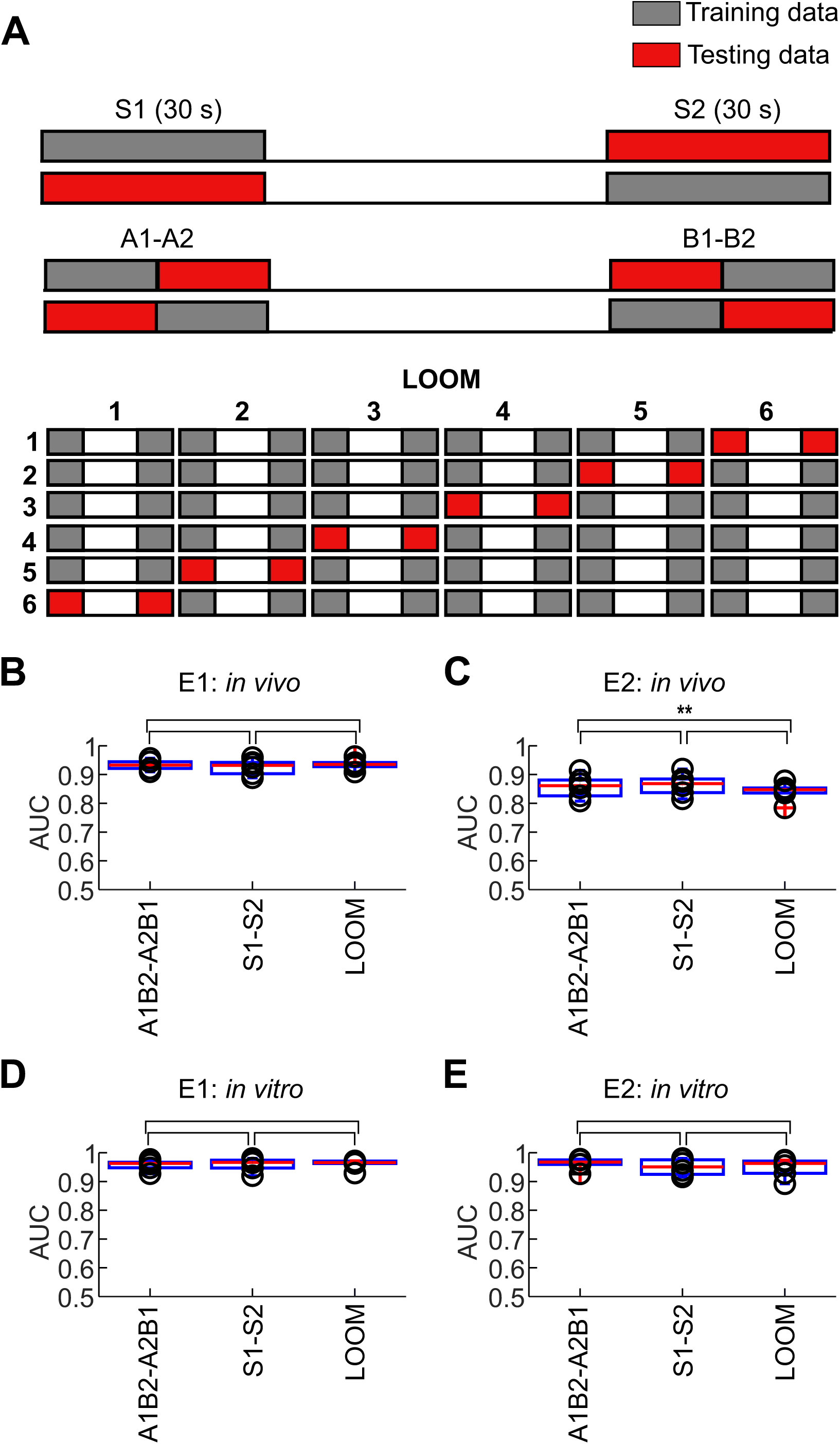
Cross-validation of MOD. (**A**) Schemes of the three cross-validation approaches “S1–S2”, “A1B2–A2B1”, and “LOOM”. Top, in the “S1–S2” scheme, training and test sets are split such that the first segment is the training set and the last segment is the test set, and vice versa. Center, in the “A1B2–A2B1” scheme, the scoring at the beginning and the scoring at the end of the recording are each split into half, and combined such that half from the beginning and half from the end are included in the training set, and the other two halves in the test set. Bottom, “LOOM” is a scheme based on a leave-one-(cell)-out approach. Data from five cells were used as the training set, and the obtained classifiers were applied to the sixth data set. This approach was repeated until the data of each cell was included exactly once in the testing set. (**B, C**) Analysis of the performance of MOD for the three cross-validation methods for six *in vivo* data sets based on two independent scorings by expert E1 (B) or expert E2 (C). The general classifier built from LOOM also provided high accuracy of detection based on E1’s manual scorings, but less so on E2’s scorings. (**D, E**) Analysis of the performance of MOD for the three cross-validation methods for six *in vitro* data sets based on two independent scorings by expert E1 (D) or expert E2 (E). Note that the AUC was consistently high for all cross-validation methods and that the general classifier worked well on the *in vitro* data sets.

### Computational efficiency

Analysis of computation times was performed on a Supermicro computer with an X9DRT mainboard, and Intel Xeon CPU (E5-2670 v2 @ 2.60GHz, Sandy Bridge, with hyper-threading enabled), using Matlab or Octave (**Table S2**). To compare the results of different lengths of the raw and training data, the computation time for each data set was normalized to a length of 30 s of training data, and 600 s of total raw data. Computing the correlation functions was the computationally most expensive part of the method (∼50 s total). Once the correlation functions were computed, solving the Wiener-Hopf equation for n = 1000 was much faster (< 0.1 s total). The application of the filter to the raw data was again computationally expensive (∼15 s total). Finally, the computation of ROC, AUC, and κ was very fast (< 0.1 s total).

When quantified per time step, filtering required *n*+1 = 1001 fused-multiply-add operations, corresponding to a computation time of ∼1 µs. This raises the intriguing possibility that a general classifier including the coefficients of the Wiener filter, the time shift δ, and the detection threshold θ_opt_ could be used for real-time filtering and detection of synaptic events during electrophysiological experiments. For such real-time applications, the best tradeoff between time shift (δ) and performance (AUC) should be set according to the results in **Fig. S3**.

### Generation of synthetic data sets

To generate ground truth data sets of synaptic activity, EPSPs were simulated in detailed passive cable models derived from reconstructed granule cells (GCs) or CA3 pyramidal neurons, using Neuron version 7.6.2 (Carnevale and Hines, 2006). Excitatory postsynaptic conductances were simulated at the dendrite, and EPSPs were measured at the soma. Excitatory postsynaptic conductances had a rise time constant *τ*_r =_ 0.2 ms, a decay time constant *τ*_d_ = 2.5 ms, and the synaptic reversal potential was E_syn_ = 0 mV. The time step of the simulations was set to 5 µs throughout. The number of segments was defined according to the “d_lambda rule”; the number of segments per section (nseg) was increased until the length of all segments was below 3.3% of the alternating current length constant at 1,000 Hz (λ_1000 Hz_; Carnevale and Hines, 2006). Simulated somatic EPSPs were analyzed using Stimfit core algorithms adapted for Mathematica (version 12.0; Guzman et al., 2014). For GCs, specific membrane resistance was set to *R*_m_ = 38,000 Ω cm^2^, specific membrane capacitance to *C*_m_ = 1 µF cm^-2^, and axial resistance to *R*_i_ = 194 Ω cm (Schmidt-Hieber et al., 2007). For CA3 pyramidal neurons, specific membrane resistance was set to *R*_m_ = 113,000 Ω cm^2^, specific membrane capacitance to *C*_m_ = 1.13 µF cm^-2^, axial resistance to *R*_i_ = 268 Ω cm (Major et al., 1994; R_m_ and C_m_ were scaled by a factor of 1.5 to account for the high density of spines in CA3 pyramidal neurons); access resistance was set to 6.9 MΩ (Vandael et al., 2020).

To test the reliability of EPSP or EPSC detection, Poisson trains of postsynaptic conductances were simulated in over 300-s time periods. Synapses were placed on the center of each dendritic branch, and activated using Neuron’s class NetStim. g_syn_ was set to 1 nS, and coefficient of peak amplitude variation was set to 0.1. Colored noise was produced by filtering of white noise with a 100-Hz first order low-pass filter, and added to the simulated traces. The signal-to-noise ratio, defined as the ratio between the event amplitude and the standard deviation of the noise, was varied between −12 and 20 dB. The MOD detection algorithm was trained on the first half (150 s) of the simulation, and then tested on the second half of the data. For GCs, current-clamp recordings of EPSPs were simulated to mimic conditions of a previously recorded *in vivo* data set (Zhang et al., 2020). For CA3 pyramidal neurons, voltage-clamp recordings of EPSCs were simulated to replicate conditions of a previous paired recording data set (Vandael et al., 2020).

To test the reliability of MOD in distinguishing frequency and amplitude changes, EPSPs with rhythmically changing frequency or amplitude were simulated (**Fig. 7**). Synaptic events were defined as alpha function templates with time constants varying in the range of 0.3 to 6 ms. The interevent intervals were drawn from an exponential distribution exp^-λ t^, in which the event rate λ was varied as appropriate.

**Fig. 6.**
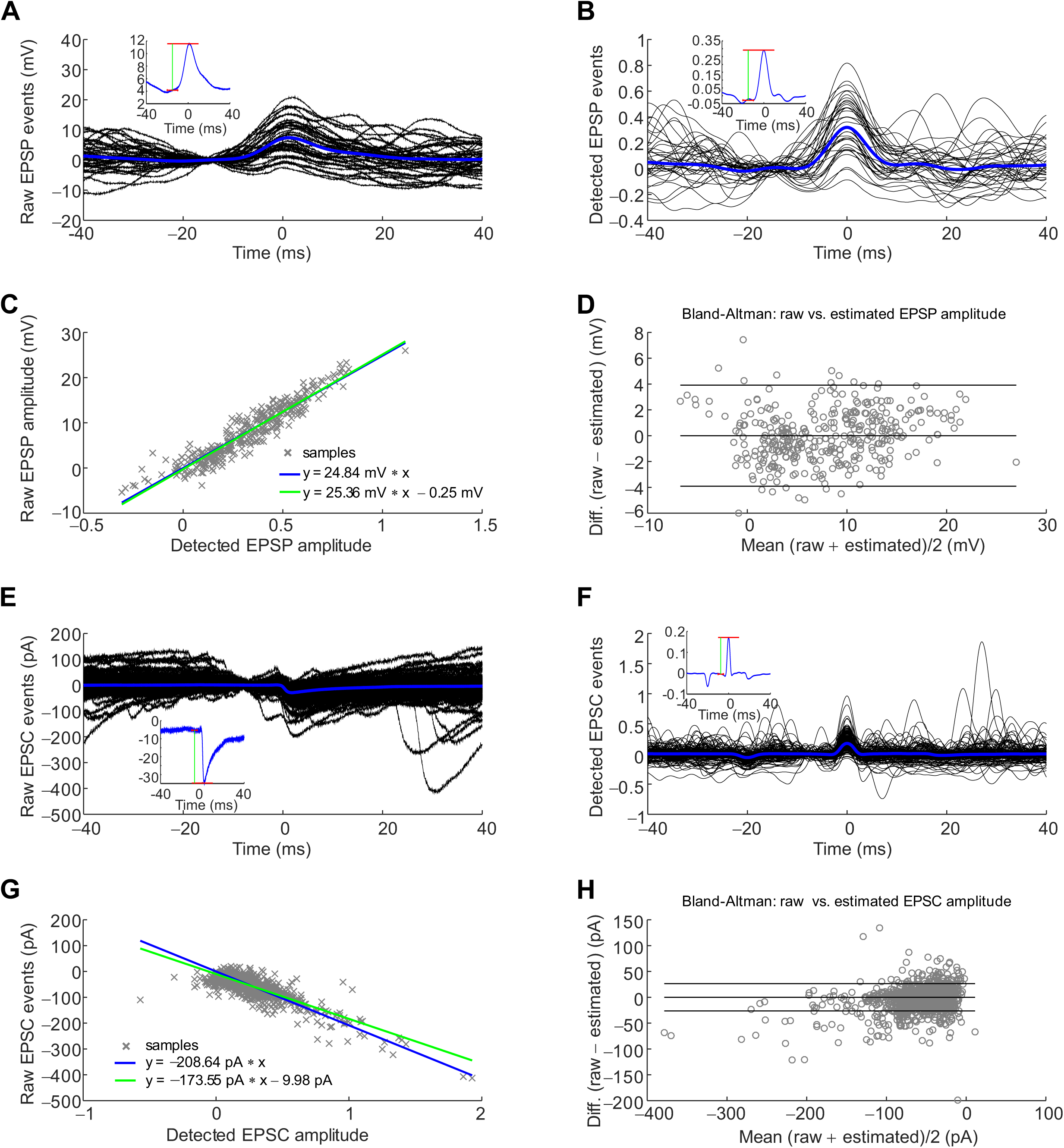
Calibrated event amplitude measurements using MOD. (**A, B**) Detection-triggered averaging of EPSPs (A) and raw detection trace epochs (B). The main panel shows superposition of individual traces (black) and average trace (blue). Insets show average traces at expanded amplitude scale. **(C)** Scatter plot of peak amplitudes of EPSP original traces versus raw detection traces. Analysis of EPSP frequency with the MOD detection method. Continuous lines represent results of linear regression (with or without offset). Slope of regression lines reveals the calibration factor. **(D)** Bland-Altman plot of difference between raw EPSP amplitude and estimated EPSP amplitude, against mean of raw and estimated EPSP amplitude. Horizontal lines indicate mean and limits of agreement (mean ± 1.96 × standard deviation). Plots in (A–D) were from *in vivo* EPSP data of a single representative cell. (**E–H**) Similar plots as (A–D), but for *in vitro* EPSC data.

**Fig. 7.**
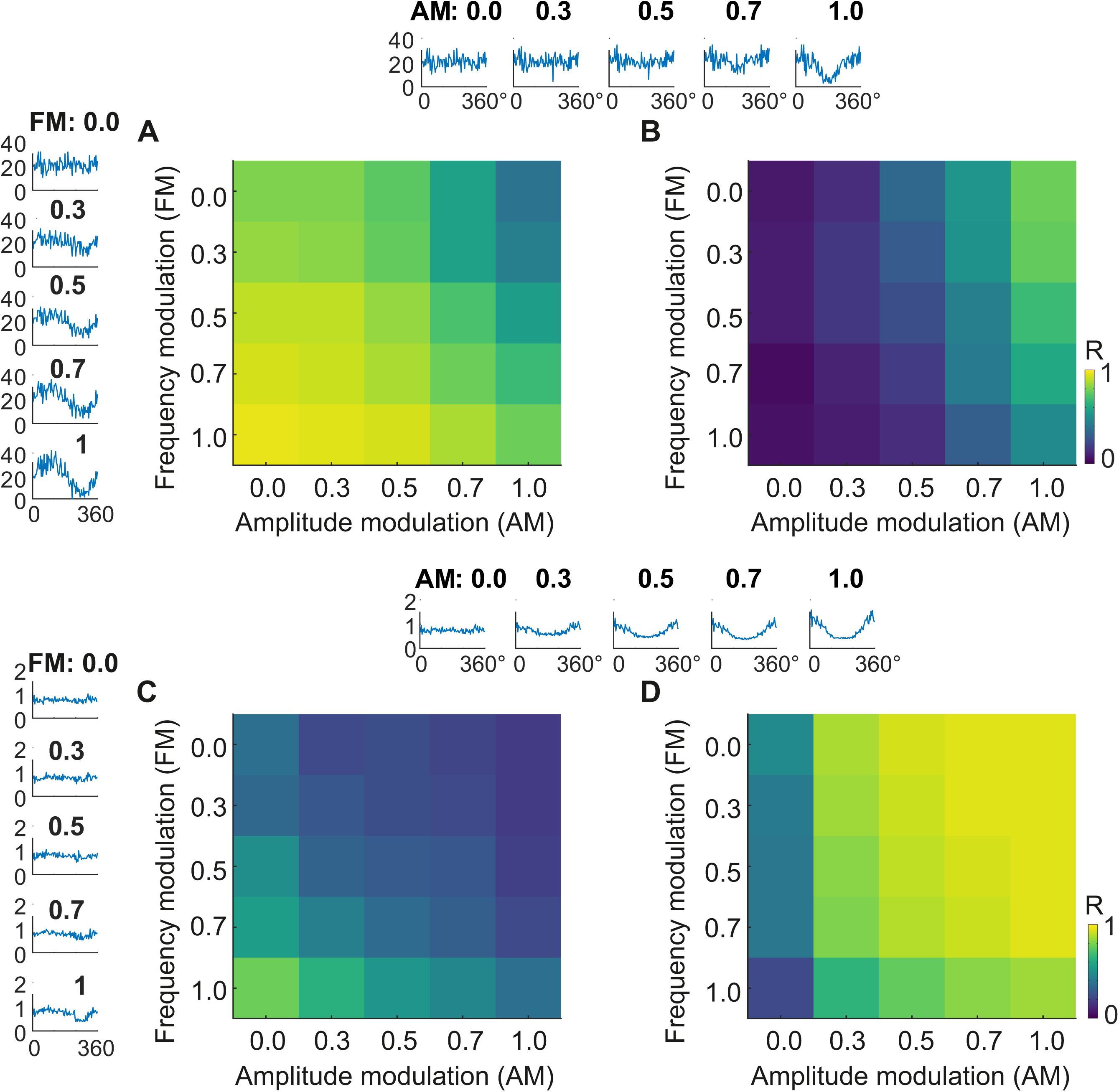
MOD analysis reliably distinguishes changes in EPSP frequency and peak amplitude. (**A, B**) Analysis of EPSP frequency with the MOD detection method. **(A)** Correlation between EPSP frequency of detected data and EPSP frequency of synthetic data. Degree of frequency modulation (FM) is plotted on ordinate, degree of amplitude modulation (AM) on abscissa; color indicates correlation between detected and simulated data; scale bar indicates correlation coefficient (R). **(B)** Correlation between EPSP frequency of detected data and EPSP amplitude of synthetic data. Left inset of traces, frequency of detected EPSPs versus location for different degrees of frequency modulation (FM, 0.0, 0.3, 0.5, 0.7, and 1.0). Top inset of traces, frequency of detected EPSPs versus location for different degrees of amplitude modulation (AM, 0.0, 0.3, 0.5, 0.7, and 1.0). (**C, D**) Similar data as shown in (A, B), but for analysis of EPSP amplitude. EPSP amplitude 1, standard deviation of colored noise 0.2; colored noise obtained by 100-Hz low-pass filtering. EPSP frequency was modulated according to a sine function, EPSP amplitude was varied according to a cosine function. Note that EPSP frequency in detected data was well correlated with EPSP frequency (A), but not amplitude in the simulated data (B). Conversely, EPSP amplitude in detected data was only minimally correlated with EPSP frequency (C), but highly correlated with EPSP amplitude in the simulated data (D). Thus, MOD analysis reliably distinguishes between changes in EPSP frequency and amplitude.

### *In vivo* recording

To further test the accuracy and efficiency of MOD, we applied the new method to biological *in vivo* data (**Fig. 3**). Experiments were carried out in strict accordance with institutional, national, and European guidelines for animal experimentation, and approved by the Bundesministerium für Bildung, Wissenschaft und Forschung of Austria (A. Haslinger, Vienna; BMWFW-66.018/0007-WF/II/3b/2014; BMWF-66.018/0010-WF/V/3b/2015; BMWFW-66.018/0020-WF/V/3b/2016).

*In vivo* recordings from the dentate gyrus GCs were performed as described previously (Pernía-Andrade and Jonas, 2014; Bittner et al., 2015; Gan et al., 2017, Zhang et al., 2020). Briefly, whole-cell patch-clamp recordings were performed in male 8- to 10-week-old head-fixed C57Bl/6J mice running on a linear belt treadmill (Royer et al., 2012; Bittner et al., 2015). The head-bar implantation was performed under anesthesia by intraperitoneal injection of 80 mg/kg body weight ketamine (Intervet) and 8 mg/kg xylazine (Graeub), followed by subcutaneous injection of lidocaine. A custom-made steel head-bar was attached to the skull, using superglue and stabilized by dental cement. After one week of recovery and another week of mild water-restriction (2 ml per day), mice were trained to run on the treadmill for 7–9 days. The day before recording, two small (∼0.5 mm in diameter) craniotomies, one for the patch electrode and one for a local field potential (LFP) electrode, were drilled at the following coordinates: 2.0 mm caudal, 1.2 mm lateral for whole-cell patch-clamp recording; 2.5 mm caudal, 1.2 mm lateral for the LFP recording. The dura was left intact, and craniotomies were covered with silicone elastomer (Kwik-Cast, World Precision Instruments). Pipettes were fabricated from borosilicate glass capillaries (1.75 mm outer diameter, 1.2 mm inner diameter). Long-taper whole-cell patch electrodes (9–12 MΩ) were filled with a solution containing: 130 mM K-gluconate, 2 mM KCl, 2 mM MgCl_2_, 2 mM Na_2_ATP, 0.3 mM Na_2_GTP, 10 mM HEPES, 18 mM sucrose, 10 or 0.1 EGTA, and 0.3% biocytin (pH adjusted to 7.3 with KOH; 310–312 mOsm). Whole-cell patch-clamp electrodes were advanced through the cortex with 500–600 mbar of pressure to prevent the electrode tip from clogging. After passing the hippocampus CA1 subfield, the pressure was reduced to 40 mbar to search for cells in the target area. After a blind whole-cell recording was obtained, series resistance was calculated by applying a test pulse (+50 mV and −10 mV) under voltage-clamp condition. Recordings were immediately discarded if series resistance exceeded 80 MΩ. All recordings were performed in the current-clamp configuration without holding current injection using a Heka EPC double amplifier. Signals were low-pass filtered at 10 kHz (Bessel) and sampled at 25 kHz with Heka Patchmaster acquisition software.

### *In vitro* recording

Paired recordings from hippocampal mossy fiber terminals and CA3 pyramidal neurons in hippocampal slices *in vitro* were performed as described previously (Bischofberger et al., 2006; Vyleta and Jonas, 2014; Vyleta et al., 2016; Vandael et al., 2020). Transverse hippocampal slices (350–400 µm thick) were prepared from 19- to 23-day-old Wistar rats of either sex (weight: 55–65 g). Mossy fiber terminals were recorded in the non-invasive cell-attached configuration, while CA3 pyramidal neurons were examined in the whole-cell recording configuration (both in voltage-clamp mode). Presynaptic and postsynaptic recording pipettes were fabricated from 2.0 mm / 1.0 mm (OD/ID) borosilicate glass tubing and had open-tip resistances of 10–20 MΩ and 3–7 MΩ respectively. For tight-seal, bouton-attached stimulation under voltage-clamp conditions, the presynaptic pipette contained a K^+^-based intracellular solution (120 mM K-gluconate, 20 mM KCl, 2 mM MgCl_2_, 2 mM Na_2_ATP, 10 mM HEPES, and 10 mM EGTA, pH adjusted to 7.3 with KOH, 297–300 mOsm). APs in mossy fiber terminals were evoked by brief voltage pulses (amplitude 800 mV, duration 0.1 ms). Postsynaptic recording pipettes were filled with an internal solution containing 130 mM K-gluconate, 20 mM KCl, 2 mM MgCl_2_, 2 mM Na_2_ATP, 10 mM HEPES, and 10 mM EGTA (pH adjusted to 7.28 with KOH, 312–315 mOsm). Postsynaptic membrane potential was set to −70 mV. Postsynaptic series resistance ranged from 3.9–14.7 MΩ, median 8.5 MΩ. After a 100-s control period, a high-frequency stimulation (HFS, 100 Hz, 1 s) was applied, followed by a 150-s test period. For MOD analysis, data before and directly after HFS were included. Data were acquired with a Multiclamp 700A amplifier, low-pass filtered at 10 kHz, and digitized at 100 kHz using a CED power1401 mkII interface (Cambridge Electronic Design, Cambridge, UK). Pulse generation and data acquisition were performed using FPulse version 3.3.3 (U. Fröbe, Physiological Institute, University of Freiburg, Germany).

### Statistics and conventions

Statistical significance was assessed using a two-sided Wilcoxon signed rank test and a Kruskal-Wallis test at the significance level (P) indicated. Box plots show lower quartile (Q1), median (horizontal red line), and upper quartile (Q3). The interquartile range (IQR = Q3–Q1) is represented as the height of the box. Whiskers extend to the most extreme data point that is no more than 1.5 × IQR from the edge of the box (Tukey style).

## Results

To detect synaptic events with high accuracy and efficiency, we developed a new method, which combines the detection power of supervised machine learning with the computational efficiency of optimal filtering. The basic work flow of the method is as follows (**Fig. 1**): First, experimenters were asked to manually score short epochs of the data containing synaptic activity (∼30 s at the beginning and ∼30 s at the end of each recording). Expert scorings were extended by short time windows of duration t_win_, resulting in a binary “manual scoring trace”. Next, a supervised machine learning-based approach, based on training and cross-validation, was used to generate an optimal filter that, when applied to the data, produced a “raw detection trace”. For an ideal detector, the raw detection trace should closely mimic the manual scoring trace. To efficiently compute the optimal filter coefficients, the Wiener-Hopf equation was applied to scoring trace and raw data (Wiener and Hopf, 1931; Wiener, 1949). As the Wiener filter had properties of a causal filter, i.e. computed data points in the raw detection trace from preceding sample points in the original data, raw detection trace and manual scoring trace were shifted by a time delay δ before comparison. Finally, the “raw detection trace” was converted into a “binary detection trace”, using a threshold value that maximized Cohen’s κ coefficient (Cohen, 1960). The optimal filter coefficients and the optimal detection threshold were then applied to both scored and unscored data (for details, see Methods). The new technique was termed MOD, which stands for **M**achine-learning and **O**ptimal-filtering-based **D**etection-procedure.

### MOD allows efficient and reliable event detection in synthetic EPSP or EPSC data sets

To assess the performance of the new method, we first tested its accuracy and efficiency on synthetic data (**Fig. 2**). Synaptic events were simulated by exponential functions generated at random intervals, and resulting traces were superimposed with colored noise. Event frequency was set to ∼10 Hz (**Fig. 2B**). Finally, the efficacy and reliability of the detection method was quantitatively probed by ROC curve analysis (Schlögl et al., 2007; Pernía-Andrade et al., 2012; Berens et al., 2018). The normalized TPR was plotted against the FPR, and area under the resulting curve (AUC) was computed (**Fig. 2C**). For an ideal detector, the AUC would approach one, whereas for complete randomness, the AUC would be as low as 0.5. As expected, the AUC approached 1 for high signal-to-noise ratio for all event frequencies and noise models tested. Remarkably, the AUC remained high even when the signal-to-noise ratios was lowered of values < 1 (**Fig. 2C, F**). Thus, MOD may be a promising technique to analyze *in vivo* intracellular recordings, in which signal-to-noise ratio is low (Lee et al., 2009; Pernía-Andrade and Jonas, 2014). To test whether the method was also suitable for analysis of EPSCs, we performed similar ground truth simulations in CA3 pyramidal neurons to mimic a previously recorded voltage-clamp data set (Vandael et al., 2020). As expected, the method was also able to accurately retrieve synaptic activity under these conditions of increased resolution.

### MOD allows efficient and reliable EPSP detection in complex *in vivo* data sets

Next, we tested the new method on biological experimental data. To rigorously test the algorithm, we applied MOD to EPSPs recorded from head-fixed mice *in vivo* during a spatial navigation task (Royer et al., 2012; Bittner et al., 2015). Patch-clamp whole-cell current-clamp recordings were made from dentate gyrus GCs (**Fig. 3A, B**). These recording conditions were highly challenging for MOD, because signal-to-noise ratio was low, synaptic events were generated at high frequency, and mechanical artifacts originating from animal movement occasionally overlaid synaptic events.

In biological data sets, the time points of generation of synaptic events are unknown. Thus, synaptic events were manually scored by two experts, one with *in vivo* recording and analysis experience (E1), and one with *in vitro* recording and analysis experience (E2). Subsequently, the MOD algorithm was trained based on these manual scorings. As for the synthetic data set, the efficacy and reliability of the method was quantitatively probed by AUC analysis (Schlögl et al., 2007; Pernía-Andrade et al., 2012; Berens et al., 2018). In total, we computed ROC curves from six *in vivo* recordings scored by two independent experts (E1, green; E2, blue; **Fig. 3C**). On average, the AUC was 0.896 ± 0.048 (mean ± standard deviation; 6 *in vivo* recordings, 12 data sets), with slightly, but significantly higher values for E1 than for E2 scoring (P = 0.0312). Thus, the new method was able to reliably detect synaptic events in *in vivo* data sets.

To benchmark the performance of the MOD method in comparison to existing methods, we analyzed the same data by template matching (TMP; Jonas et al., 1993; Clements and Bekkers, 1997; Chadderton et al., 2004) and deconvolution (DEC; Pernía-Andrade et al., 2012; **Fig. 3D, E**). For the TMP method, an alpha function template was used, whereas for the DEC method an instantaneous exponentially decaying template was employed (Pernía-Andrade et al., 2012). In both cases, templates with time constants ranging from 1–50 ms were tested, and the best performing AUC result was used to provide a maximally rigorous comparison with MOD. Comparison of the AUC values of all three methods revealed that the MOD method was statistically superior to the template-fit and the deconvolution method for both scorings (P = 0.00376; **Fig. 3D, E**). On average, the degree of improvement, quantified from the ratio (1−AUC)_MOD_ / (1−AUC)_TMP_ or (1−AUC)_MOD_ / (1−AUC)_DEC_, was 260% and 314% for E1 and 208% and 271% for E2. Thus, MOD is the method of choice for the analysis of subthreshold synaptic activity in *in vivo* data sets.

### MOD improves EPSC detection in *in vitro* data sets

Next, we tested whether the new MOD method might also convey advantages for the analysis of synaptic activity *in vitro*. In particular, we wondered whether MOD might improve detection under conditions, in which event frequency was increased. To test this, we made paired recordings from presynaptic mossy fiber terminals and postsynaptic CA3 pyramidal cells (**Fig. 4A**). Mossy fiber terminals were stimulated non-invasively with a 1-s, 100-Hz train of pulses in the cell-attached mode, while adjacent CA3 pyramidal neurons were recorded in the whole-cell voltage-clamp configuration (Bischofberger et al., 2006; Vyleta et al., 2016; Vandael et al., 2020). These recording conditions were also challenging for MOD, because EPSC frequency was enhanced during and after high-frequency stimulation (**Fig. 4B**).

As for the *in vivo* data set, short epochs of data in six *in vitro* data sets were manually scored by the two experts. Subsequently, manual scoring was used to train the MOD algorithm. Finally, the detection efficiency and reliability was probed by AUC analysis (**Fig. 4C**). On average, the AUC was 0.960 ± 0.005 (6 *in vitro* mossy fiber bouton–CA3 pyramidal neuron recordings, 12 data sets), similar for scoring by E1 and E2 (P = 0.84). Benchmarking against previously published techniques, template-fit and deconvolution, revealed that MOD showed a significantly higher performance (P = 0.00376; **Fig. 4D, E**). On average, the degree of improvement, quantified from the ratio (1−AUC)_MOD_ / (1−AUC)_TMP_ or (1−AUC)_MOD_ / (1-AUC)_DEC_, was 201% and 265% for E1 and 290% and 519% for E2. Thus, MOD was not only highly advantageous for the analysis of subthreshold EPSPs *in vivo* data sets, but also led to marked improvement in the efficacy and accuracy of detection of EPSCs in *in vitro* data sets.

### Optimal parameter settings for the MOD algorithm

We then examined how the settings of various parameters affected the accuracy and efficiency of MOD analysis. First, we varied the time window for manual scoring (t_win_; **Fig. S1**). For both *in vivo* and *in vitro* data sets, the highest AUC was obtained for t_win_ = 3 ms or 4 ms. Second, we varied the filter order, and plotted AUC against filter order over sample frequency, i.e. corresponding filter duration (**Fig. S2**). For both *in vivo* and *in vitro* data sets, a filter duration of at least 20 ms was required for accurate detection. Third, we explored how the temporal shift δ between manual scoring trace and raw detection trace would affect the reliability of MOD analysis (**Fig. S3**). For *in vivo* data sets, the optimal value of the time shift δ was ∼20 ms. Thus, the method worked best when it relied on experimental data that occurred ∼20 ms in the past before a given time point (**Fig. S3A, B**). However, for the *in vitro* data sets, the optimum value of time shift δ was reached at ∼5 ms (**Fig. S3C, D**). These differences correspond to differences in the total duration of the synaptic events, which are slow EPSPs under *in vivo* conditions versus faster EPSCs under *in vitro* conditions. Finally, we analyzed how the power of the detection algorithm was affected when the duration of the scored epochs was reduced (**Fig. S4**). When total duration of the scored epochs was reduced from 60 s to 20 s or 10 s, the distribution of the AUC values changed only minimally, for both *in vivo* and *in vitro* data sets. Thus, for our experimental data, a total duration of scored epochs of 10 s seemed to be sufficient to ensure reliable analysis. However, when the total duration of the scored epochs was reduced to values of 1–5 s, a component with AUC values < 0.8 appeared in the distributions. This indicates a higher probability that the trained filter would fail if the training data was very short. Thus, consistent results from MOD analysis cannot be guaranteed for such short scoring periods. In conclusion, these results indicate that reliable MOD analysis requires a t_win_ of 3–4 ms, a filter duration of at least 20 ms, a time shift δ that approximately matches the time course of the underlying synaptic events, and a minimal scoring period of ∼10 s.

### Cross-validation of MOD suggests generalization to unscored data

A potential problem with machine-learning approaches is their tendency to over-fit the data (Kevin and Murphy, 2012). If training and test set are the same or overlap, the classifier may work extremely well on the trained data, but may fail to generalize to unscored data. To test whether the MOD method generalizes to new data, we applied three different cross-validation schemes (**Fig. 5**). In the “S1–S2” scheme, training and test sets were split such that the first segment (S1) was included in the training set and the last segment (S2) in the test set, and vice versa (**Fig. 5A**, top). In the “A1B2–A2B1” scheme, we concatenated the first half of S1 (A1) and the second half of S2 (B2) as training data, and the other two halves as test data (A2B1; **Fig. 5A**, center). Such a scheme may account for possible non-stationarities during the recording period. Both approaches (S1–S2 and A1B2–A2B1) used classifiers that were built from individual cells. Additionally, we used a cross-validation scheme based on a LOOM (**Fig. 5A**, bottom). Manually scored data sets from five cells were combined to form the training set, and the obtained classifier was applied to the data from the sixth cell. The procedure was repeated until each cell was included exactly once in the test set (**Fig. 5A**, bottom). This cross-validation scheme allowed us to test the performance of a “general classifier” when applied to unscored data sets.

We then compared the results from three different cross-validation schemes for *in vivo* and *in vitro* data sets scored by both experts (**Fig. 5B–E**). For all cross-validation schemes, the AUC values were close to 0.9, indicating that the MOD detection algorithm generalized well to unscored data. Unexpectedly, the differences between the cross-validation schemes were only minimal. For example, the AUC values derived from LOOM cross-validation analysis were not significantly different from those computed by A1B2–A2B1 or S1–S2, suggesting that it may be possible to use a general classifier for future recordings obtained under similar conditions (**Fig. 5D, E**). This property of the MOD algorithm is potentially useful for large-scale analysis of synaptic activity, because it reduces the amount of manual scoring work.

For a subset of experiments, particularly data recorded *in vivo*, the AUC was slightly higher for A1B2–A2B1 than for S1–S2 (**Fig. 5B**). This may be due to nonstationarities in recording conditions (e.g. input resistance or access resistance) or alterations in the behavioral state of the mice. Thus, expert scoring only at the beginning or the end of the recording may not be sufficient to provide adequate analysis of data over the entire recording period. Moreover, for the *in vivo* data analyzed by expert E2, the AUC values for LOOM cross-validation were slightly lower than those for the A1B2–A2B1 scheme (P = 0.00376; **Fig. 5C**). This may suggest that a classifier built from the scoring of expert E1 (who has extensive experience in the analysis of *in vivo* data) generalizes better to previously unscored data sets than a classifier based on the scoring of expert E2 (who has experience in the analysis of *in vitro*, but not *in vivo* data). Thus, for building general classifiers, it may be important to base the analysis on the scoring of experts with a high level of specific expertise. In conclusion, the MOD method can be used for automated large-scale analysis of synaptic activity, as required for the understanding of information encoding in neuronal populations.

### MOD allows accurate detection of both time points and peak amplitudes

Our results indicate that MOD allows us to accurately retrieve the time points of synaptic events in both *in vivo* and *in vitro* data sets. However, the method does not directly reveal information about peak amplitudes of synaptic events. We reasoned that it might be possible to infer peak amplitudes of synaptic events from peak amplitudes of the raw detection trace after appropriate calibration. To determine the calibration factor, we performed event-triggered averaging for the *in vivo* data set, in which the EPSP amplitude was measured under current-clamp conditions (Zhang et al., 2020; **Fig. 6**). Both original traces (Fig. **6A**) and raw detection traces (**Fig. 6B**) were aligned by the time points of the detected events, and averaged across traces. The amplitude calibration factor was then determined as the ratio of peak amplitudes of the two signals. To further corroborate linearity of the amplitude calibration, we plotted the peak amplitude of EPSP original traces against those of the raw detection trace. Scatter plot analysis revealed a linear dependency of original and detection trace data (**Fig. 6C**). Bland-Altman analysis further revealed that the two quantities are in excellent agreement, corroborating the reliability of our estimates (**Fig. 6D**). Similar results were obtained for *in vitro* data sets, in which the EPSC amplitude was measured under voltage-clamp conditions (Vandael et al 2020; **Fig. 6E–H**). Taken together, these results indicate that MOD allows us to precisely determine both time points and peak amplitudes of synaptic events.

Finally, we tested whether MOD was able to distinguish changes in event frequency and peak amplitude when the two quantities were changing in an overlapping, time-dependent manner (**Fig. 7**). We simulated synaptic events with variable exponential time course, varied events frequency (F) according to a sine function, and amplitude (A) according to a cosine function. F-F analysis, in which the correlation between frequency of detected events and frequency of original data was examined, revealed that frequency modulation was accurately retrieved (**Fig. 7A**). Furthermore, F-A analysis, in which the correlation between frequency modulation of detected events and amplitude modulation of the original data was tested, indicated that frequency analysis was only minimally perturbed by concomitant amplitude changes (**Fig. 7B**). In addition, A-F and A-A correlation analysis showed that amplitude changes were accurately retrieved, whereas overlaying frequency changes only minimally perturbed the results (**Fig. 7C, D**). In conclusion, these results indicate that MOD is able to dissect temporal changes in event frequency and peak amplitude, as required for a mechanistic analysis of single-neuron computations under *in vivo* conditions.

## Discussion

Precise quantitative analysis of subthreshold synaptic activity in neurons *in vivo* and *in vitro* is essential for the understanding of both the biophysical properties of synaptic transmission and the synaptic mechanisms of neural coding. However, detecting EPSPs and EPSCs remains challenging, especially under *in vivo* conditions. In this paper, we report a new method for detecting synaptic events based on machine learning and optimal filtering, termed MOD. AUC analysis indicated that the new method is superior to template-fit and deconvolution methods. Under typical recording conditions, the new method can lead to an up to 3-fold increase in detection performance, as quantified by the (1–AUC)^-1^ metric. Thus, the new method may be useful to generate new insights into the mechanisms of single-neuron computations and the cellular mechanisms of neuronal coding.

### Advantages of MOD

The new algorithm has several advantages. First, and foremost, it offers by far the highest area under the ROC curve. Thus, for any previously specified FPR, a markedly smaller false negative rate can be obtained. Similarly, for any given false negative rate, a much smaller FPR can be achieved. Second, the threshold selection based on Cohen’s κ coefficient is fully automated and objective. Thus, the new method lacks free parameters, in contrast to template-fit methods, deconvolution algorithms, and Bayesian methods, which all rely on assumptions about template kinetics and detection thresholds (Jonas et al., 1993; Clements and Bekkers, 1994; Pernía-Andrade et al., 2012; Merel et al., 2016). Third, cross-validation used in the MOD algorithm provides quantitative information about the expected detection errors of unscored data. The new algorithm has obvious advantages for the analysis of subthreshold activity *in vivo*, but it is also useful for detection of synaptic events *in vitro*. While low-frequency miniature synaptic activity can be reliably detected by standard methods, high-frequency activity, for example during asynchronous transmitter release (Hefft and Jonas, 2005), is more difficult to analyze.

Our results provide a proof-of-principle that such high-frequency activity is reliably detected with the new method.

### Potential limitations of MOD

An apparent disadvantage of the new method is that manual scoring of a subset of data is required to train the algorithm to obtain the optimal filter coefficients. However, manual scoring of relatively short time periods, ∼10 s (**Fig. S4**), is sufficient to optimize the detection. Furthermore, our cross-validation analysis (LOOM; **Fig. 5**) suggested that in data sets obtained under similar experimental conditions, the same filter coefficients may be used across cells. Thus, manual scoring may be required only in a subset of representative recordings, and could be applied more widely on unscored recordings.

Another potential disadvantage is that the MOD analysis is computationally more demanding than template-fit (Jonas et al., 1993; Clements and Bekkers, 1997; Chadderton et al., 2004) or deconvolution (Pernía-Andrade et al., 2012). However, this primarily applies to the training phase, in which the optimal filter coefficients are determined and optimal threshold and time shift values are found. In contrast, the filtering phase is comparatively fast, only requiring computation times of ∼1 µs per time step (**Table S2**). Thus, MOD is a computationally efficient method, especially under conditions where large unscored portions of data need to be analyzed or general classifiers can be applied to unscored cells.

Finally, the performance of MOD may depend on the quality of the expert scoring. Our two expert scorings were similar but not identical. While systematic differences in the choice of the fiducial point were corrected by MOD via the time shift δ (**Fig. S3**), annotation jitter or inconsistent judgement of individual events may pose a problem. Interestingly, for the *in vivo* data set, the AUC values were consistently higher for expert E1 (having extensive experience in the analysis of *in vivo* data) than for expert E2 (having experience with *in vitro*, but not *in vivo* data). Thus, MOD may be able to learn better from a user with specific expertise. Furthermore, for the *in vivo* data set, LOOM, A1B2–A2B1, and S1–S2 cross-validation gave almost identical results for E1 scoring, whereas LOOM produced significantly lower AUC values for E2 scoring. Thus, MOD may generalize better to unscored cells if trained by a user with specific expertise. These aspects should be taken into account, especially when attempts are made to build general classifiers.

### Future applications

We used the MOD technique for the analysis of EPSPs *in vivo* in awake, behaving mice running on a linear belt and of EPSCs *in vitro* before and after high-frequency stimulation. However, several additional applications are conceivable. The MOD method may be useful to analyze neuronal coding in invertebrates, in which analogue signaling is more prevalent than in mammals (Debanne et al., 2012). Furthermore, the MOD algorithm could be utilized for the analysis of intracellularly recorded spikes or spikelets (Epsztein et al., 2010). Finally, the computational efficacy of the filtering step of MOD may allow experimenters to run the detection in real time. Although the current implementation involves non-causal filtering (Eq. 8), a causal filter could be generated by introducing an additional delay between recording and detection. Our analysis of AUC as a function of time shift δ between manual scoring trace and raw detection trace indicates saturation at δ ≈ 10 ms (**Fig. S3**). Thus, the delay could be set to relatively short values without compromising the power of the detection method. Therefore, MOD could be used in closed-loop *in vivo* experiments, in which real-time detection of synaptic events is coupled to either electrical stimuli or optogenetic manipulation of defined cell populations (Scanziani and Häusser, 2009; Grosenick et al., 2015). Thus, MOD not only may become important for data analysis, but also could be integrated into cutting-edge experimental methodology.

## Acknowledgements

This project has received funding from the European Research Council (ERC) under the European Union’s Horizon 2020 research and innovation programme (grant agreement number 692692 to P.J.) and the Fond zur Förderung der Wissenschaftlichen Forschung (Z 312-B27, Wittgenstein award to P.J.). We thank Drs. Jozsef Csicsvari, Christoph Lampert, and Federico Stella for critically reading previous manuscript versions. We also thank Florian Marr for technical assistance, Eleftheria Kralli-Beller for manuscript editing, and the Scientific Service Units of IST Austria for efficient support.

## Author contributions

PJ conceived the project, AS and PJ developed algorithms, XZ performed *in vivo* experiments and analyzed data, DV performed *in vitro* experiments, and AS, XZ, and PJ wrote the paper. All authors jointly revised the paper.

## Code availability

The core functions of the newly developed MOD method will be publicly available under GPLv3. All other codes used to generate the presented results will be available from the corresponding author upon reasonable request.

## Data availability

The data sets used for benchmarking will be available from the corresponding author upon reasonable request.

## Figure Legends

**Fig. S1.**
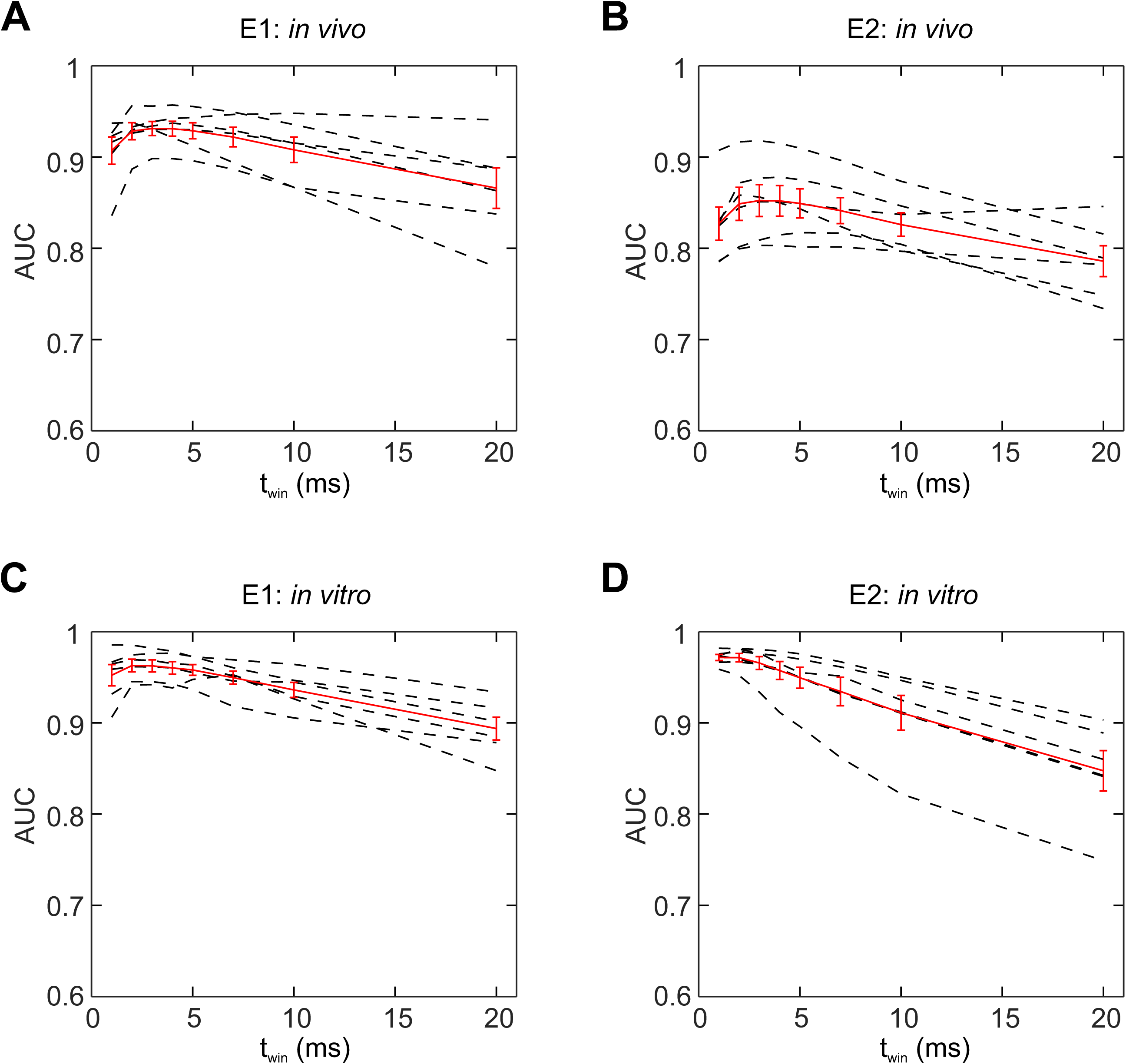
The optimal time window for manual scoring (*t*_win_). Analysis of the dependence of AUC on the duration of the manual scoring expansion window (*t*_win_). Data were obtained from *in vivo* recordings (**A, B**) or *in vitro* recordings (**C, D**) and scored by either expert E1 (**A, C**) or expert E2 (**B, D**). Note that the optimum *t*_win_ value was 3–4 ms under all conditions. Red curve, mean ± SEM; dashed lines, data from individual experiments. The optimum value of *t*_win_ defines the temporal resolution of the entire method, including manual scoring and automatic detection. Results shown were computed using the cross-validation scheme “A1B2–A2B1”.

**Fig. S2.**
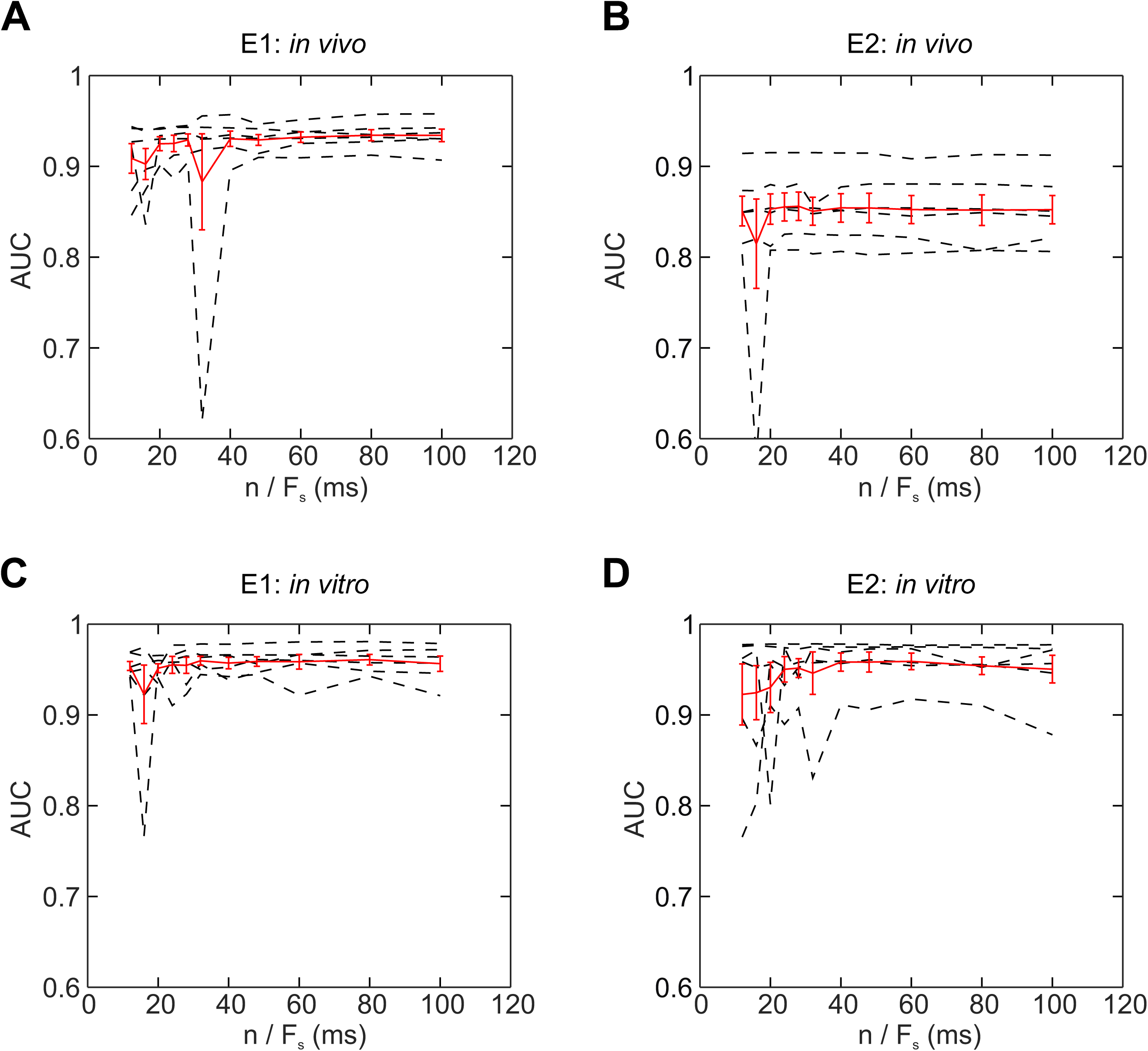
Effects of the order of the Wiener filter. Analysis of the dependence of AUC on the order of the Wiener filter. Filter length was obtained as the ratio n / F_s_, where n indicates number of filter coefficients in Eqs. 1 and 2 and F_s_ represents sampling frequency. Data were obtained from *in vivo* recordings (**A, B**) or *in vitro* recordings (**C, D**) and scored by either expert E1 (**A, C**) or expert E2 (**B, D**). Red curve, mean ± SEM; dashed lines, data from individual experiments. Results shown were computed using the cross-validation scheme “A1B2–A2B1” with *t*_win_ = 4 ms.

**Fig. S3.**
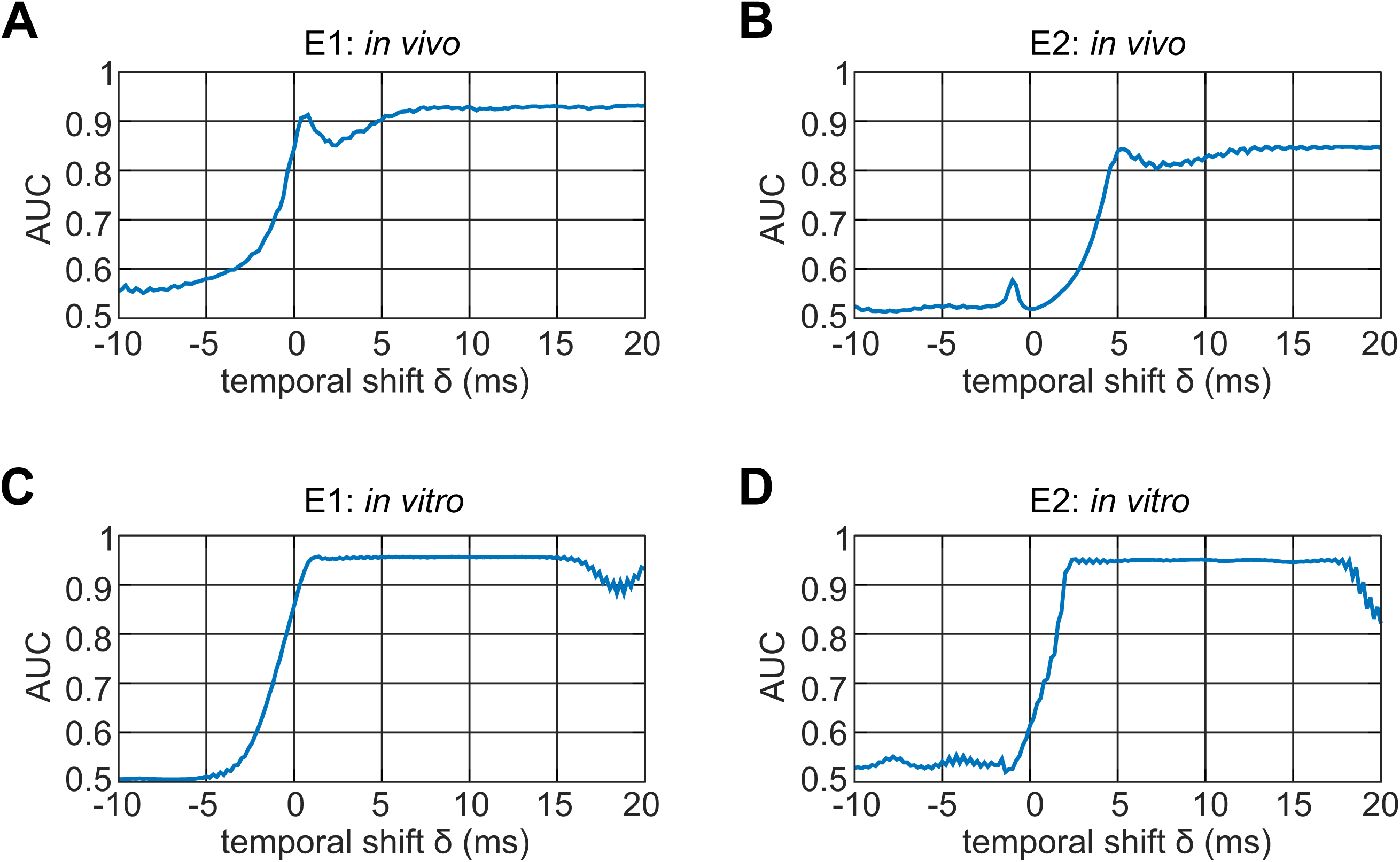
The optimal time shift δ of the Wiener filter. Analysis of the dependence of AUC on the time shift δ, where δ = 0 corresponds to the time points of manual scoring. Data were obtained from *in vivo* recordings (**A, B**) or *in vitro* recordings (**C, D**) and scored by either expert E1 (**A, C**) or expert E2 (**B, D**). Note that the optimum time shift was close to 20 ms for the *in vivo* data sets, but close to 5 ms for the *in vitro* data sets. This is likely to be caused by the slower time course of EPSPs in the *in vivo* data in comparison to the faster time course of the EPSCs in the *in vitro* data. Also note that delay values were slightly different between expert scorings (E1 versus E2), because experts marked synaptic events at slightly different time points. Results shown were computed using the general classifier.

**Fig. S4.**
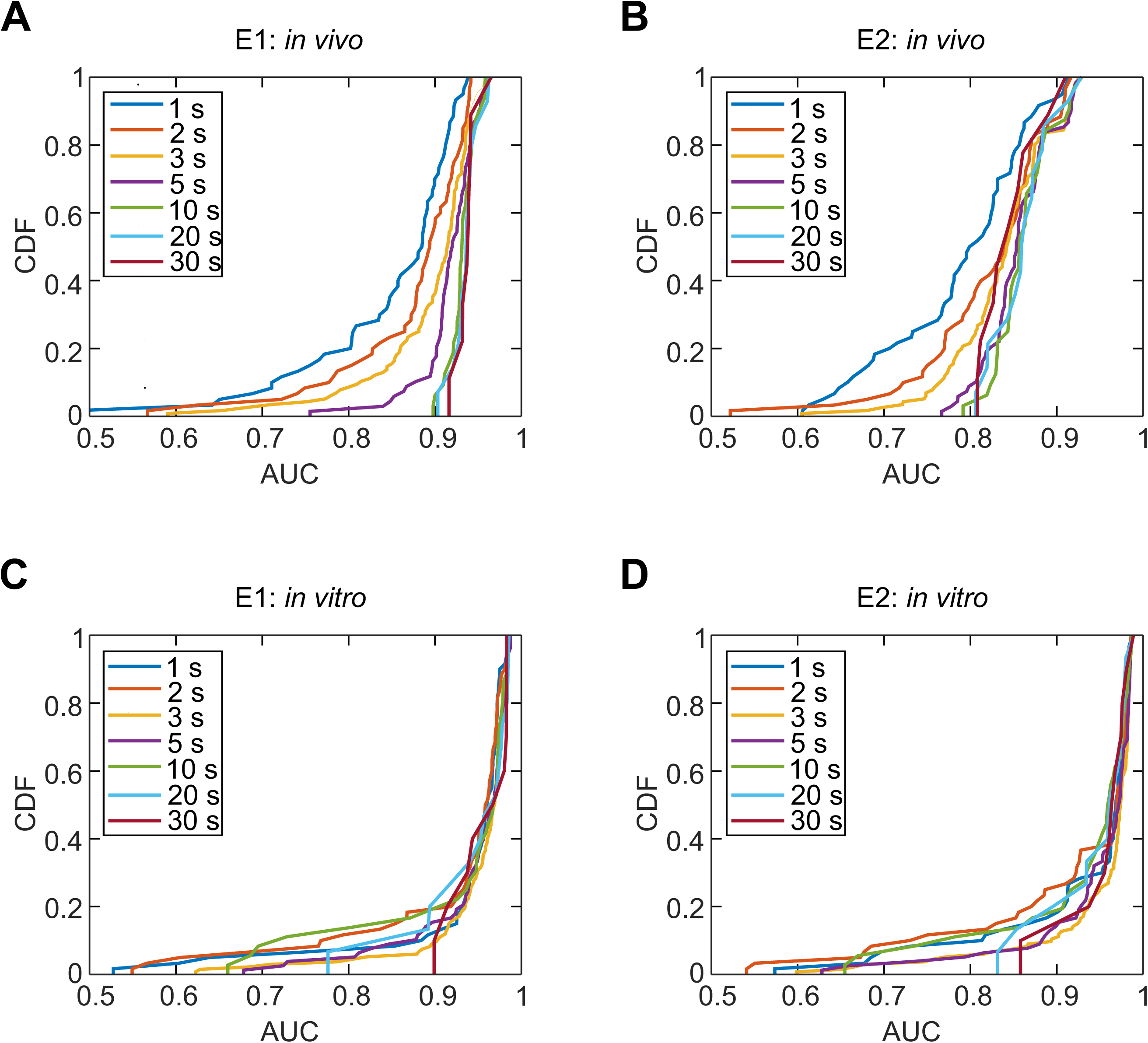
The minimal length of scored traces required for reliable detection. Cumulative distribution of AUC values for different duration of the scoring period used for training of MOD. Data were obtained from *in vivo* recordings (**A, B**) or *in vitro* recordings (**C, D**) and scored by either expert E1 (**A, C**) or expert E2 (**B, D**). Note that AUC values were consistently above 0.8 for scoring periods of ≥ 10 s. In contrast, a fraction of analysis data points showed AUC values of < 0.8 for shorter scoring periods of 1–5 s. Thus, MOD analysis based on shorter scoring periods is possible, but occasionally unreliable.

**Fig. S5.**
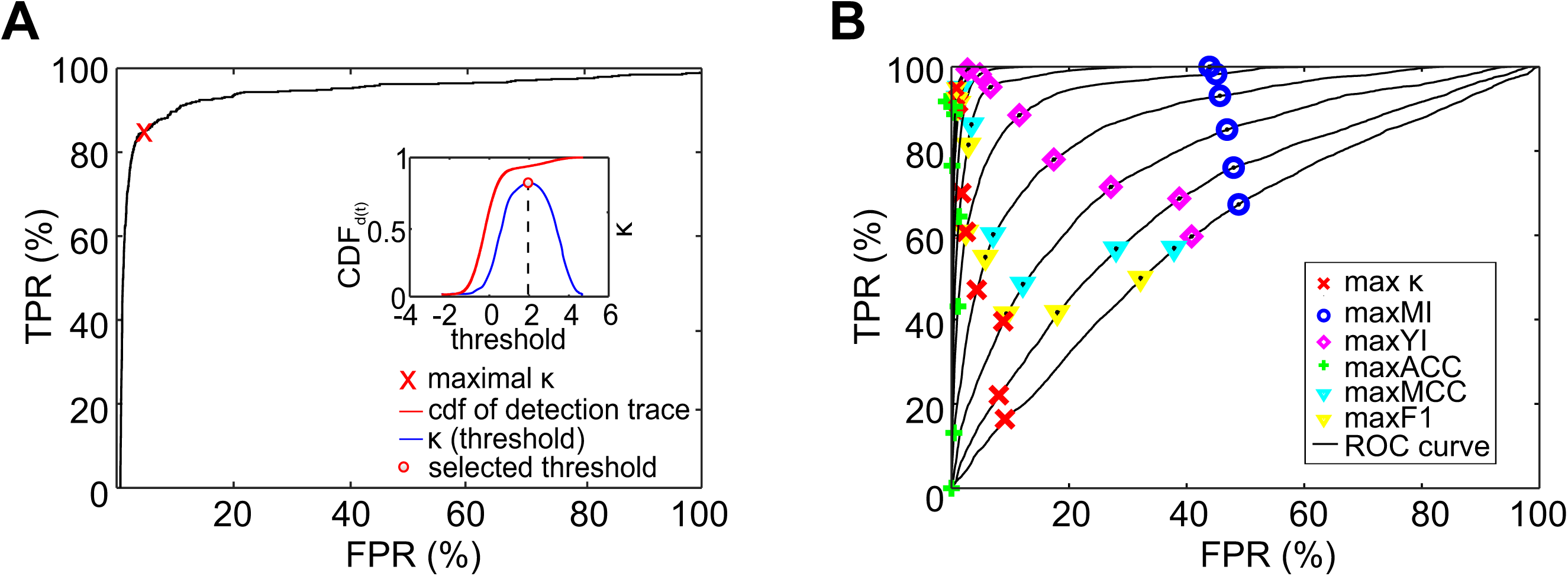
Comparison of different criteria for determining detection threshold. **(A)** Plot of TPR against FPR for a synthetic data set (ROC curve). The signal-to-noise ratio was 3, i.e. 9.5 dB in this simulation. The AUC was 0.970. The “x” in the ROC curve indicates the point on the ROC curve that corresponds to maximal κ, implying the optimal detection threshold. Inset shows the dependence of Cohen’s κ on the threshold, peak value of κ, and cumulative distribution of data points in the detection trace. **(B)** ROC curves for simulated data. The signal-to-noise ratio SNR = 20*log_10_(A_peak/rms_noise) was varied from −52 dB to 26 dB in steps of 7.8 dB. Note that larger signal-to-noise ratios yield ROC curves with a larger AUC. The maxima of different performance metrics (Cohen’s κ, mutual information (MI), Youden index (YI), accuracy (ACC), Matthews correlation coefficients (MCC), and F1) are depicted.

**Table S1.** Properties of manually scored data sets. *In vivo* data set was obtained by making whole-cell current-clamp recordings of EPSPs in dentate gyrus GCs near the resting potential in head-fixed mice running on a linear belt (for details, see Methods). *In vitro* data set was obtained using presynaptic cell-attached recording from hippocampal mossy fiber terminals and simultaneous postsynaptic whole-cell voltage-clamp recordings of EPSCs from CA3 pyramidal neurons in acute hippocampal slices (for details, see Methods).

| Parameter | *in vivo* data set | *in vitro* data set | Additional information |
| --- | --- | --- | --- |
| Total scoring duration | 71.0 ± 10.1 s | 73.7 ± 8.4 s |  |
| Total number of scored events | 2100 ± 683 | 811 ± 305 |  |
| Average event frequency | 30.0 ± 9.3 Hz | 11.4 ± 5.6 Hz |  |
| Number of scored cells | 6 | 6 |  |
| Number of scorings | 12 | 12 | Expert E1 with *in vivo* background, expert E2 with *in vitro* background |

**Table S2.** Computation time for individuals steps of MOD. The results were computed on a Supermicro computer with an X9DRT mainboard, and Intel Xeon CPU (E5-2670 v2 @ 2.60GHz, Sandy Bridge, with hyper-threading enabled). The machine was running under the operating system “Debian(9.6)/Stretch”, and the computations were done with Octave 4.4.0 on a single CPU core. Computation time is wall clock time in seconds. Twelve data sets were analyzed, and the cross-validation scheme (A1B2–A2B1) was used, thus 24 results were computed. The computational time of each data set was normalized to a length of 30 s of training data, and 600 s of total raw data.

| | Computation time<br>(mean ± SD) | Reference | Conditions |
| --- | --- | --- | --- |
| Computing auto-correlation functions | 14.14 ± 0.32 s | Eq. 6a | normalized to 30 s of training data |
| Computing cross-correlation functions | 35.33 ± 0.79 s | Eq. 6a | normalized to 30 s of training data |
| Solving Wiener-Hopf equation | 0.028 ± 0.001 s | Eq. 5 | |
| Raw data filtering | 14.59 ± 0.26 s | Eqs. 2 and 8 | normalized to 600 s of raw data |
| Computing ROC, AUC, and $\kappa$ | 0.073 ± 0.044 s | Fig. 1c;<br>Fig. 2c, d | normalized to 600 s of raw data |

